# Structural characterization of an RNA bound antiterminator protein reveal a successive binding mode

**DOI:** 10.1101/2020.09.27.315978

**Authors:** James L. Walshe, Karishma Patel, Sandro F. Ataide

## Abstract

In bacteria, rho-independent termination occurs through the formation of an intrinsic RNA terminator loop, which disrupts the RNA polymerase elongation complex, resulting in dissociation from the DNA template. ANTAR domains are a class of antiterminator that bind and stabilise mutually exclusive dual hexaloop motifs within the nascent RNA chain to prevent terminator loop formation. We have determined the structures of the dimeric ANTAR domain protein EutV, from *E. faecialis,* both in the absence of and in complex with the dual hexaloop RNA target. The structures illustrate the conformational changes that occur upon RNA binding and reveal the molecular interactions between the ANTAR domains and RNA are restricted to a single hexaloop of the motif. Our findings thereby redefine the minimal ANTAR domain binding motif to a single hexaloop and revise the current model for ANTAR-mediated antitermination. These insights will inform and facilitate the discovery of novel ANTAR domain RNA targets.

## INTRODUCTION

Transcription can be broadly categorized into three highly regulated processes: initiation, elongation and termination. Termination results in the irreversible dissociation of the RNA polymerase complex (RNAP) from DNA and prevents unintended gene expression, interference from antisense transcripts and provides the cell with a mechanism to rapidly respond to changes in the extracellular environment (1–4). In bacteria, termination occurs either through the action of the Rho RNA helicase or the formation of intrinsic termination loops (T-loops) (1, 4). T-loops account for approximately 80% of RNA ends in *E. coli* (5) however, their abundance varies across bacterial species (6). Intrinsic termination occurs when the RNAP complex stalls on a poly-uridine tract long enough for the preceding GC rich sequence to fold into a T-loop (Figure 1A) within the RNA exit tunnel thus destabilizing the RNAP complex leading to transcriptional termination (7). T-loops provide a means to demarcate gene boundaries. However, as their formation is passive, bacteria require mechanisms to allow the RNAP to bypass T-loops or prevent T-loops from folding in the first place: a process known as antitermination (2, 8)

**Figure 1.**
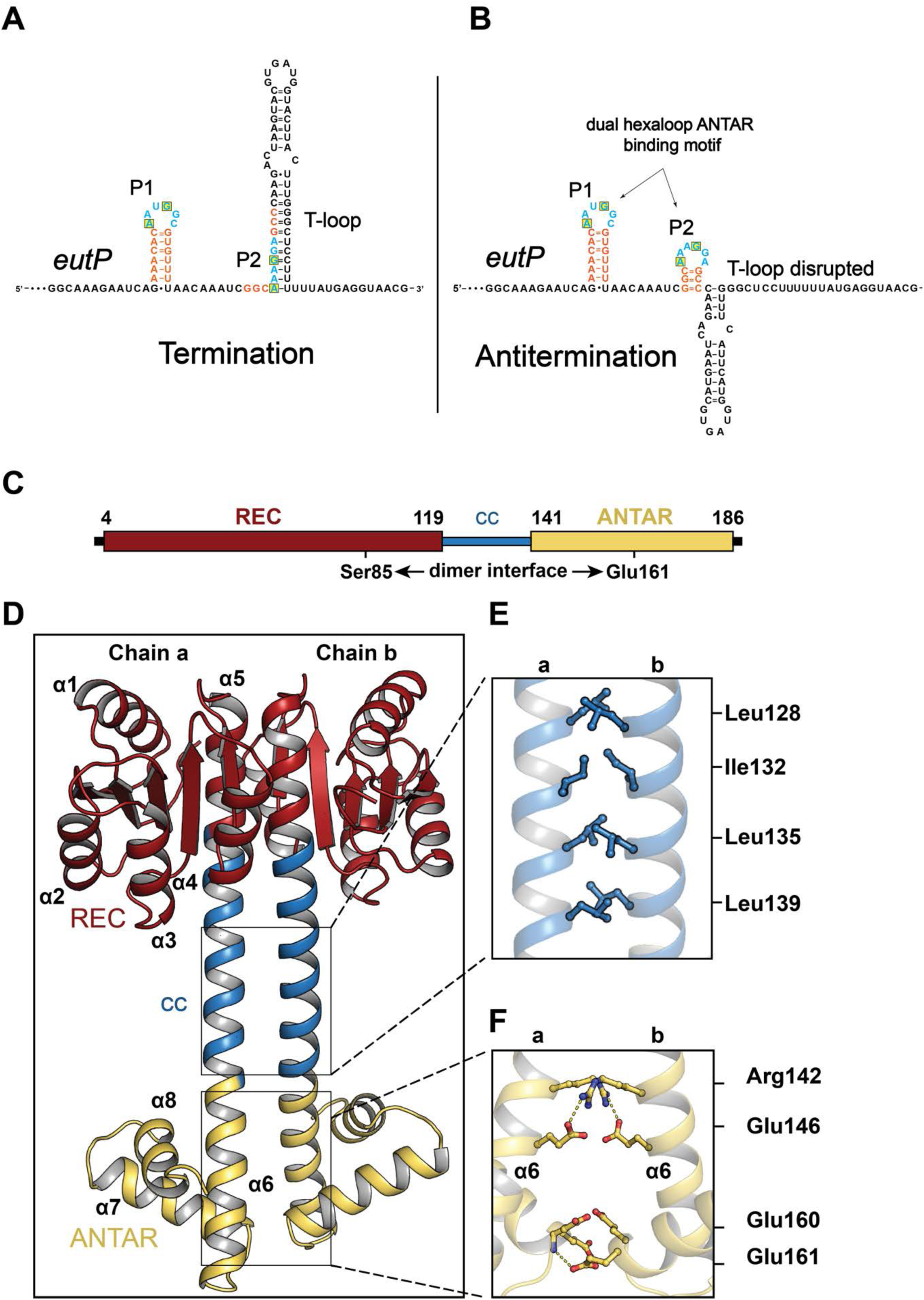
ANTAR binding motif upstream of *eutP* gene and structure of *E. faecalis* antitermination protein EutV. (A-B) Primary sequence of 5’ UTR of the *eutP* gene from *E.faecalis eut* operon. Bases involved in stem formation are shown in orange, hexaloop bases in blue and conserved bases at position 1 and 4 of the hexaloops highlighted in a yellow box. (A) Schematic showing the folding of the intrinsic terminator. Only the P1 hexaloop of the ANTAR binding motif is folded with P2 hexaloop embedded within the intrinsic terminator loop. (B) Alternative folding for the intrinsic terminator region into a dual hexaloop motif promoted by the binding of dimeric EutV. Both P1 and P2 hexaloop’s are folded (24,33,34). (C) Schematic representation of dual domain nature of EutV with the receiver (REC) domain and the ANTAR domain shown in red and yellow respectively. (D) 2.1 Å resolution crystal structure of EutV captured in a dimeric state highlighting the (E) hydrophobic packing of the coiled-coil residues and (F) hydrogen bonding between the ANTAR domains.

Passive antitermination mechanisms that affect the formation of T-loops include the action of RNA- binding antiterminator proteins, stalled ribosomes, uncharged tRNAs or small molecules via riboswitches (9–11). Antiterminator proteins bind specific sequences and/or structural elements in the nascent chain RNA in order to prevent T-loop formation. HutP, from *B. subtilis*, directly binds T-loop sequence repeats to regulate expression of the *hut* operon (12), while the BlgG/SacY/LicT/GlcT family of antiterminator from *B. subtilis* and *E. coli* bind a structured RNA element that precludes the T-loop from forming (13–18). These antiterminator proteins contain vastly different RNA binding domains and display distinct RNA binding mechanisms (Supplementary Figure 1), highlighting the need for individual study of novel antitermination proteins.

The AmiR and NasR transcription antitermination regulator (ANTAR) domain was first characterised in AmiR, a positive regulator of the *P. aeruginosa* amidase operon, (19–23) and was subsequently identified in the well-studied antiterminator protein NasR (23). ANTAR domains are a novel output domain for two-component signalling (TCS) pathways that, unlike the majority of output domains, bind to RNA instead of DNA (24–26). Transduction pathways using TCS exist in bacteria, some archaea, plants and lower eukaryotes (27–29), and allow for the rapid conversion of an external signal to an intracellular response. The ANTAR domain consists of a three-helical bundle of approximately 60 amino acids, with a conserved aromatic residue exposed to the cavity formed by the three-helix structure (23). Proteins containing ANTAR domains are typically modular and include coiled-coil or low complexity domains for dimerization, along with a vast array of sensor or receiving domains such as c**G**MP-specific phosphodiesterases, **a**denylyl cyclases and **F**hlA (GAF) and Per-Arnt-Sim (PAS) domains (Supplementary Figure S2 A). The diversity of these accompanying domains is emphasised by the four known structures of ANTAR domain proteins (Supplementary Figure S2 B-E)(19,30–32) and highlight the large range of bacterial metabolic and regulatory processes that ANTAR domains govern. Current structural studies have focused on the unbound state of ANTAR domains and despite recent efforts to determine ANTAR residues involved in RNA binding through mutagenesis (33) the molecular details of the interaction between an ANTAR domain and its target RNA remains unknown.

Recently, the consensus RNA binding motif for ANTAR domain proteins was characterised in the *E. faecalis eut* operon, and found to consist of a dual hexaloop motif, with positions 1 and 4 of the hexaloops conserved as adenine and guanine bases respectively (Figure 1B) (25). These motifs are capable of folding within all four T-loops of the *eut* operon (Supplementary Figure 3A-B) with the second hexaloop overlapping with the stem of the T-loop, providing a discernible means for antitermination upon ANTAR domain binding (Figure 1A-B) (25). Furthermore, dual hexaloop motifs have since been identified in other characterised ANTAR domain systems (25, 26). Regulation of T-loop formation in the *eut* operon of *E. faecalis* is under the control of the EutW/EutV TCS pathway. Ethanolamine (EA) stimulates the histidine kinase EutW (component 1) to phosphorylate EutV (component 2) which promotes homodimerization of the protein (Supplementary Figure S3 A). Dimeric EutV acts as an antiterminator of RNAP by binding these dual hexaloop motifs within the nascent mRNA chain inhibiting T-loop formation and thereby preventing transcription termination (25).

ANTAR domains are implicated in the regulation of a wide variety of bacterial processes through both TCS systems or direct coupling to protein sensor domains (22,32,34–37)(Supplementary Figure S2 A). Additionally, an ANTAR domain involved in the regulation of light sensitive processes, through attachment to a light-oxygen-voltage (LOV) receptor, was recently identified in *N. multipartite* (Supplementary Figure S2 C) (31), paving the way for potential optoribogenetic approaches. Given the expanding interest in ANTAR domains and the plethora of bacterial processes they have been implicated in the regulation of, understanding the molecular detail of the interactions between ANTAR domains and their target dual RNA hexaloops is crucial for our understanding of these systems and the potential development of therapeutics to disrupt them. Here we report the first crystal structures of EutV both in the absence of and in complex with the dual hexaloop RNA target. These discoveries allow us to propose a revised model for ANTAR mediated antitermination whereby an ANTAR domain dimer contacts each hexaloop of the dual hexaloop motif successively in order to prevent termination in eubacteria. We redefine the minimal ANTAR domain binding motif to a single hexaloop which will help facilitate the discovery of novel ANTAR domain RNA targets.

## RESULTS

### Structure of RNA-free EutV antitermination protein

EutV possesses one of the most common domains architectures found amongst all ANTAR domain- containing proteins annotated in the Pfam database (Supplementary Figure 2A) (49), consisting of an N-terminal phosphor-sensitive REC domain (aa 4-119) coupled to a C-terminal ANTAR domain (aa 141-186) via a coiled-coil domain (Figure 1C). To ascertain the structural arrangements of the ANTAR and REC domains of EutV relative to one another, crystals of recombinantly expressed full-length EutV were produced and diffraction data collected to 2.1 Å resolution (Supplementary Table 1, PDB ID: 6WSH). Within the crystal, EutV showed a symmetric dimer “dumbbell” conformation with 48 residues contributing to 1870 A^2^ of buried surface area and extensive hydrophobic and ionic interactions between residues Ser85 and Tyr164 of each chain (Figure 1D). The REC domain of each chain dimerised through a common ‘α4-β5-α5’ mode (50) forming one end of the dumbbell with the ANTAR domains forming the other end. The handle of the dumbbell consists of a coiled-coil that includes residues Leu128, Ile132, Leu135 and Leu139 that form the majority of hydrophobic interactions within a perfect heptad repeat (Figure 1E). The interaction between the two ANTAR domains is stabilised by mutual hydrogen bonds between Arg142 to Glu146 (2.9 Å and 3.0 Å), and the amine group of chain a Glu161 to Glu160 chain b (3.1 Å) (Figure 1F).

EutV is monomeric in solution (25) however SEC-MALS data indicates that EutV can form higher order species, likely a dimer (Supplementary Figure S4) in the absence of phosphorylation. The dimeric state within the crystal structure was unexpected given the recombinant EutV used for crystallography was not phosphorylated nor supplemented with phosphomimetics. However, rather than acting as a definitive switch, phosphorylation of REC domains is known to shift the equilibrium of dynamically sampled states (51–53). In this way, during crystallisation and with an increased concentration of EutV, the dimeric state was likely preferentially favoured.

### Structure of RNA bound EutV antiterminator protein

#### Model building the RNA bound EutV structure

To determine the molecular interactions of the ANTAR domain of EutV with the dual hexaloop RNA substrate, EutV in the presence of the phosphomimetic beryllium fluoride was crystallised in complex with a 51-nt RNA corresponding to the 5’ UTR of *eutP* (Figure 2A), that contains both the P1 and P2 hexaloops (25), to 3.8 Å resolution (Supplementary Table 1, PDB ID: 6WW6, Figure 2D). Molecular replacement was performed using the dimeric RNA-free EutV model to estimate initial phases and allowed for the modelling of two asymmetric EutV chains. Unambiguous electron density for a single RNA hexaloop was present at the ANTAR domain of each EutV chain (Supplementary Figure S5). Modelling of the entire 51-nt substrate, with a single hexaloop contacting each ANTAR domain, was not possible given the relative orientation of the hairpins bound at each ANTAR domain. The 5’ ends of each hexaloop were over 70 Å apart with only seven RNA bases unmodeled (Supplementary Figure S6 A-B). It became apparent that each of the modelled single hexaloops within the asymmetric unit (ASU) were from different RNA molecules. The dual hexaloop RNA molecule was bridging between two asymmetric units within the crystal lattice with each hexaloop contacting a symmetry-related ANTAR domain of EutV in the neighbouring ASUs (Supplementary Figure S7 A-B). To confirm this, the web based RNAComposer software program (26) was used to generate an idealised three-dimensional structure of the 51-nt RNA. These coordinates, when manually fitted to the positive electron density, were the ideal length to stretch between ANTAR domains from neighbouring ASU in both directions within the crystal lattice (Supplementary Figure S7 C-D).

**Figure 2.**
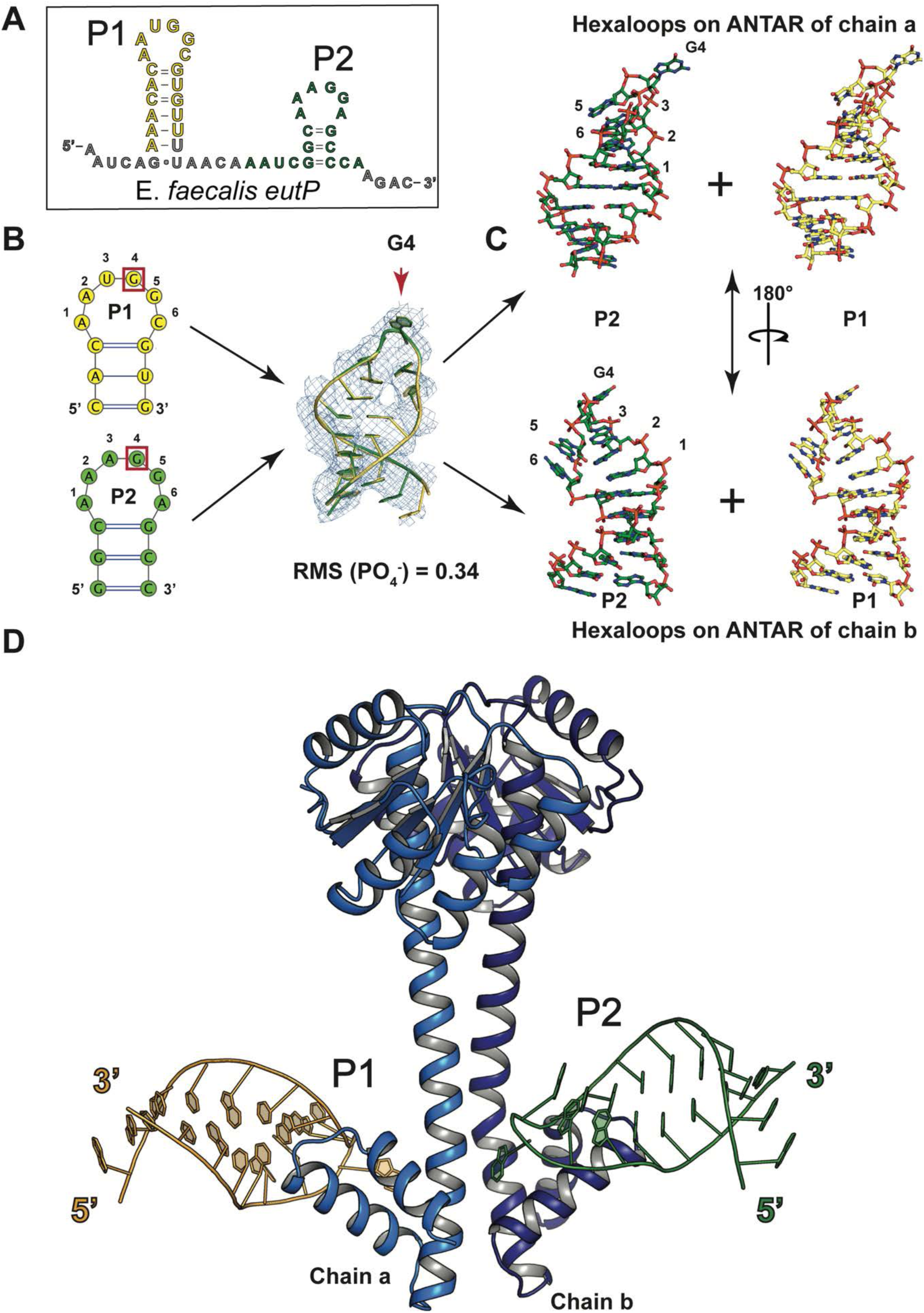
Structure of EutV bound to the antitermination RNA motif. (A) *EutP* dual hexaloop sequence from *E. faecalis eut* operon within the crystal. P1 and P2 hairpins shown in yellow and green respectively. (B) Cartoon representation showing an alignment of the ribose and phosphate backbone of both the P1 and P2 hexaloop from each ANTAR domain of the EutV dimer highlighting the near identical position of the bases. 2F_o_-F_c_ map contoured to at 1 σ (C) Stick representation of each for hexaloops described in (B). (D) Crystal structure of EutV bound to e*utP* dual hexaloop motif.

Because each RNA hexaloop (P1 or P2) has a different primary sequence (Figure 2A-B), and each hairpin of the same RNA molecule contacts the symmetry related ANTAR domain in neighbouring ASUs (Supplementary Figure S7 D), the electron density for the RNA hairpin at either ANTAR domain within a single ASU represents a combination of the sequences of the P1 and P2 hexaloops (Figure 2B). Within the crystal, whenever a P1 hairpin makes contact with an ANTAR domain within a discrete ASU, the P2 hairpin of the same RNA molecule must contact the symmetry-related ANTAR domain in a neighbouring ASU. Therefore, a model was built by placing both the P1 and P2 hairpins into the density at each ANTAR domain of the dimer, and refinement then performed with a fixed 50% occupancy for all nucleotides (Extended data Table 1). Despite consisting of different sequences, the P1 and P2 hairpins contacting the ANTAR domains from separate RNA molecules refined to near identical conformations (RMSD = 0.34 Å for the phosphate and ribose backbone) (Figure 2B, Supplementary Figure S8). In total, each ASU contains an asymmetric dimer of EutV and two RNA hexaloops from different RNA molecules.

In both hairpins, a single base of the hexaloop flips outward to interact with EutV. The flipped nucleotide is either in position 3 or 4 of the hexaloop, depending on the direction the hairpin is modelled (Figure 2B-C, Supplementary Figure S8). The antitermination motif includes a conserved guanosine at positions 4 (G4) of each hexaloop that are obligatory for efficient EutV mediated antitermination *in vivo* (25). Therefore, the flipped base was defined as G4 and used to orientate the direction of the RNA hexaloops on each ANTAR domain. Given this contact is the only base specific interaction between the protein and RNA hairpins, and this base is in an identical location in both modelled hairpins (Figure 2B), it can be described with confidence despite the averaging effect applied to the electron density. For clarity, only one of the two possible RNA hexaloops at each ANTAR domain will be described in the rest of the manuscript and are labelled P1 and P2 (Figure 2D).

#### Protein:RNA interactions and binding sites

Despite the limited resolution (3.8 Å) successful modelling of EutV residues that interact with the RNA hairpins was achieved through use of the recently developed *ISOLDE* package designed for building high-quality macromolecular models into low to medium resolution experimental maps (Supplementary Figure S9) (46). The interaction between dimeric EutV and the antitermination RNA hexaloops is restricted to the ANTAR domain of each chain, consistent with published gel retardation assays performed using truncated EutV constructs (25). Two putative RNA binding sites with asymmetric protein:RNA interactions were identified, one at each ANTAR domain of the EutV dimer (Figure 3). The different binding modes are due to the large difference between the kink in the coiled-coil region of chain a and that of chain b, which is clearly noticeable when each chain is compared to the RNA-free EutV dimer (Supplementary Figure S10 A-B). As a result, chain a and b make more interactions with the P2 RNA hexaloop than P1. Both sites bind a single hexaloop with the conserved G4 of each loop flipping outward to form π–π stacking interactions with Tyr164 of the ANTAR domain (Figure 3A-C). Additionally, the hydroxy group of Tyr164 makes a hydrogen bond to the phosphate group of the base in position 3 of the hexaloop (Figure 3C). The G4 of P2 is coordinated by hydrogen bonds to Arg142, Lys149 and Glu160 of chain b, however, Arg142 from chain a is unable to make the same contact with P1 (Figure 3, compare A and C). Intriguingly, residues Lys143 and Lys147 from chain a contact the P2 RNA, through hydrogen bonds to the phosphate and ribose respectively, representing the only interactions between an ANTAR domain and the RNA hexaloop located on the opposite protein chain (Figure 3B). The prominent nature of the kink in the coiled-coil of chain b prevents the reciprocal interactions between residues Lys143 and Lys147 of chain b existing with P1 (Figure 3D).

**Figure 3.**
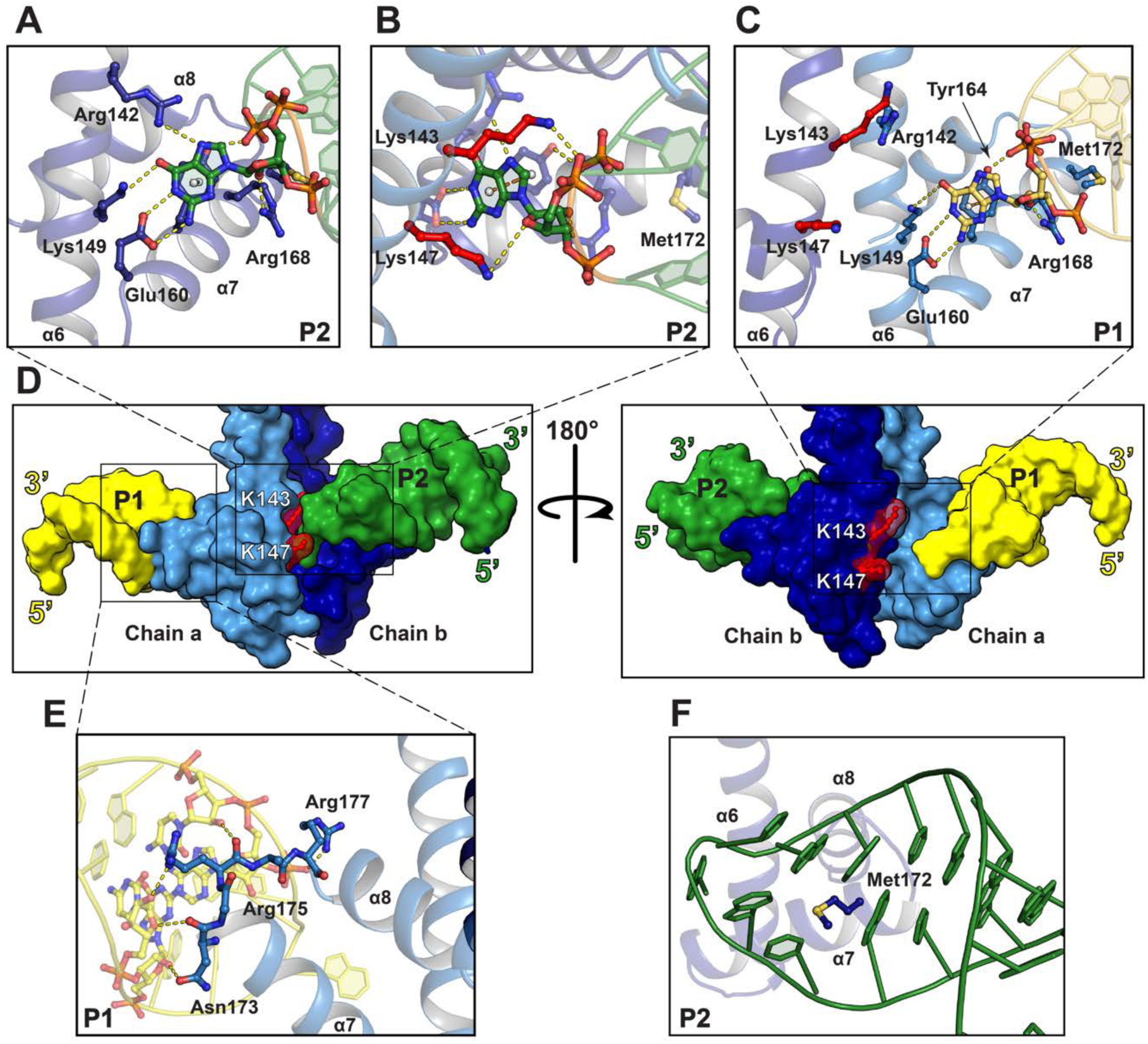
Interactions between ANTAR domain and hexaloop. (A) Interactions between the ANTAR domain of chain b and the P2 hexaloop. (B) Interaction between chain a ANTAR with the P2 hexaloop. (C) Interactions between chain a ANTAR domain and P1 RNA hexaloop. (D) Surface representation of the RNA bound EutV structure with Lys143 and L147 highlighted in red (E) Interaction between loop α7-α8 loop and P1 hexaloop (F) Met172 of chain b ANTAR domain positions P2. Chain a and b shown in light and dark blue respectively. Hydrogen bonds shown as yellow dashes.

On both chain a and chain b, Arg168 is well positioned to hydrogen bond to the ribose hydroxyl group of the RNA (Figure 3 A-C). Additionally, the residues of the α7-α8 loop make similar contacts with the RNA backbone on either chain. Asn173 and Arg175 hydrogen bond with hydroxyl groups of the ribose sugars of the bases that form the stem of the hexaloop, and Arg177 makes a hydrogen bond to the phosphate backbone of the base at position 2 of the hexaloop (Figure 3E). As the interactions are limited to the closing three base pairs of the RNA stem, they provide a molecular understanding for the obligatory nature of these stems in *in vivo* antitermination, independent of their sequence (25). Unexpectedly, given its hydrophobic nature, Met172 is positioned in the middle of the hexaloop of the RNA hairpin and may act as a hydrophobic plug to position the hexaloop on α7 within the ANTAR domain and flip the G4 base out of the hexaloop (Figure 3F). Met172 may also form potential S- H/π interactions with bases within the hexaloop (54).

### Alanine Mutagenesis

The majority of ANTAR residues implicated in RNA binding are highly conserved (Figure 3A-E, Figure 4A). To validate the role of these residues, six full length EutV constructs containing alanine mutations were generated. The folded-states of the mutant constructs were confirmed to be identical to wildtype by circular dichroism and one-dimensional NMR (Supplementary Figure S11 A-B), and their RNA binding ability was assessed using surface plasmon resonance (SPR) (Figure 3B and Supplementary Figure S12). Single alanine mutations to the residues involved in base specific interactions with the flipped G4 base resulted in a respective 60- and 65-fold reduction in binding affinity for E160A and Y164A (Figure 3B, Supplementary Figure S12 A-C). The third G4 coordinating mutant, R142A, showed a less dramatic decrease in binding with only an 8-fold decrease and the double mutant, K143A/K147A, had a similar modest effect on RNA binding (Figure 4B, Supplementary Figure S12 D-E). Drastically, the N173A/R175A/R177A triple mutation and the M172A mutation completely abolished binding to the RNA (Figure 4B, Supplementary Figure S12 F-G). The decrease in binding seen across all mutant constructs highlights the significance of the RNA interacting residues identified within the crystal structure and provides a molecular rationale for the findings of the recent mutagenesis studies performed on the ANTAR domain protein AmiR from *P.* aeruginosa (33).

**Figure 4.**
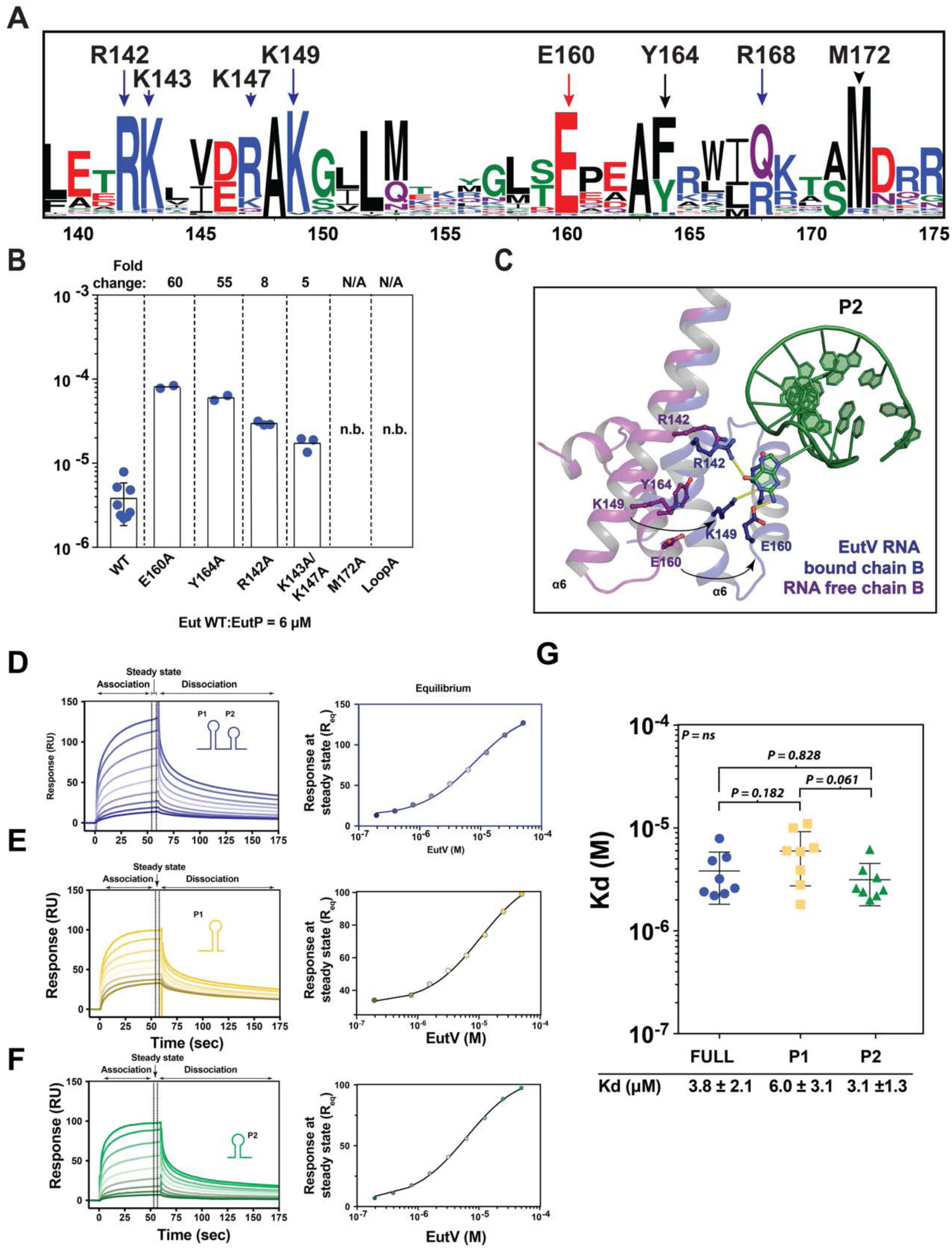
EutV binding to single RNA hexaloops. (A) Multiple sequence alignment 2332 ANTAR domain proteins containing a N-terminal REC domain and C-terminal ANTAR domain shown as a WebLogo (64). (B) Binding affinity of EutV mutants to eutP RNA as determined by surface plasmon resonance. (C) Movements of residues involved in RNA binding between RNA-free and RNA bound EutV structures. (D-F) Representative normalised SPR sensorgrams of EutV binding to the *eutP* RNA and the P1 and P2 hexaloops. Association and dissociation regions are shown above panel and apply for all sensorgrams. Sensorgrams showing increasing EutV concentrations (0, 0.4, 0.8, 1.5, 3.125, 6.25, 12.5, 25, 50, 100 μM). Representative dose response plot of the interaction of EutV with immobilised RNA (as shown in corresponding left panel) at equilibrium fitted to a one-site Langmuir isotherm. (G) Average experimental *K_D_* values from eight SPR experiments are 3.8 ± 2.1, 6.0 ± 3.1 and 3.1 ±1.3 μM for, *eutP*, P1 and P2 hexaloop respectively. Error bars are standard deviations of eight separate runs. The affinity of EutV to each of the three RNAs was compared between using an analysis of variance (ANOVA), with the individual means being compared using a Tukey’s HSD test to maintain an overall 5% error rate. The overall p value was 0.06 and none of the means were different at the 5% level.

### EutV binding to P1 and P2 RNA

The RNA-bound EutV structure clearly indicated an asymmetric binding mode and shows that a EutV dimer cannot bind to both hairpins of the dual hexaloop motif at the same time. The structure of P1 and P2 hexaloops are close to identical with an RMSD of 0.34 Å and both hairpins contain the signature flipped G at position 4 on the hexaloop (Figure 3 A-C). To elicit if EutV displayed a binding preference for either the P1 or P2 hairpin, we determined the binding affinity of EutV to *eutP* or the individual P1 and P2 RNA hairpins using SPR (Figure 4 D-E). No significant difference (P=0.061) was observed between EutV binding to the *eutP,* P1 and P2 RNAs (6.0 ± 3.1 and 3.1 ±1.3 μM respectively) (Figure 4 G). This was consistent with the crystal packing arrangement where the P1 or P2 hexaloops contact both EutV chains throughout the crystal (Supplementary Figure S7 D). This raises questions as to the previously identified obligatory role of both P1 and P2 hairpins in *in vivo* EutV mediated antitermination (25).

## DISCUSSION

In the presence of ethanolamine, EutW phosphorylates EutV on a conserved Asp54 residue in the REC domain, promoting homodimerization (25). The dimeric RNA-free structure of EutV resembles the crystal structure of the ANTAR domain protein AmiR from *P.* aeruginosa (Supplementary Figure S13 A-B). Like EutV, AmiR is a positive regulator of gene expression through a C-terminal ANTAR domain. However, AmiR itself is regulated by the direct interaction of AmiC with its N-terminal pseudo-REC domain that lacks residues required to accept phosphorylation from a TCS kinase (Supplementary Figure 13 A). Thus, it has remained unclear how phosphorylation triggers dimerization of EutV.

Within the EutV structure, the conserved active site residues in the N-terminal REC domain are in a “phospho-activated” state, as indicated by their position relative to the to the beryllium fluoride activated CheY/CheZ complex (Figure 5A). This indicates the dimeric structure of EutV that we observed is representative of the biologically active phosphorylated dimer required for efficient antitermination (25,55–59). Despite its dimeric structure, EutV shares sequence similarly (37%) and an identical domain architecture with the antitermination protein Rv1626 from *M. tuberculous* (Supplementary Figure S14) (30, 36). Like EutV, in the absence of phosphorylation Rv1626 exists as a monomer in solution and was crystallised in a monomeric state (Figure 5C-D) but is thought to form an extended dimeric structure upon phosphorylation (30). The differences between the two crystal structures arise from the extended nature of the α5-α6 helix in EutV relative to monomeric Rv1626 (Figure 5C, Supplementary Figure S14). The active site residues in the REC domain of Rv1626 are in the non-phosphorylated state (Figure 5B-C), highlighted by an “outward” facing Tyr111 (Tyr101 in EutV) that sterically inhibits a α4-β5-α5 dimer interface from forming as present in the EutV structure (Figure 5C). Given the sequence similarity between the two proteins, it is likely that the monomeric state of EutV seen in solution (Supplementary Figure S4) will adopt a similar compact conformation to Rv1626, representing the inactive state for antitermination regulation. Comparison of the monomeric Rv1626 and the EutV dimer allows us to model the extension of the α5-α6 helices that a monomeric EutV would be required to undergo, upon phosphorylation of Asp54 (25, 35), to form an extended state capable of dimerization through the coiled-coil domain (Figure 5 B-D).

**Figure 5.**
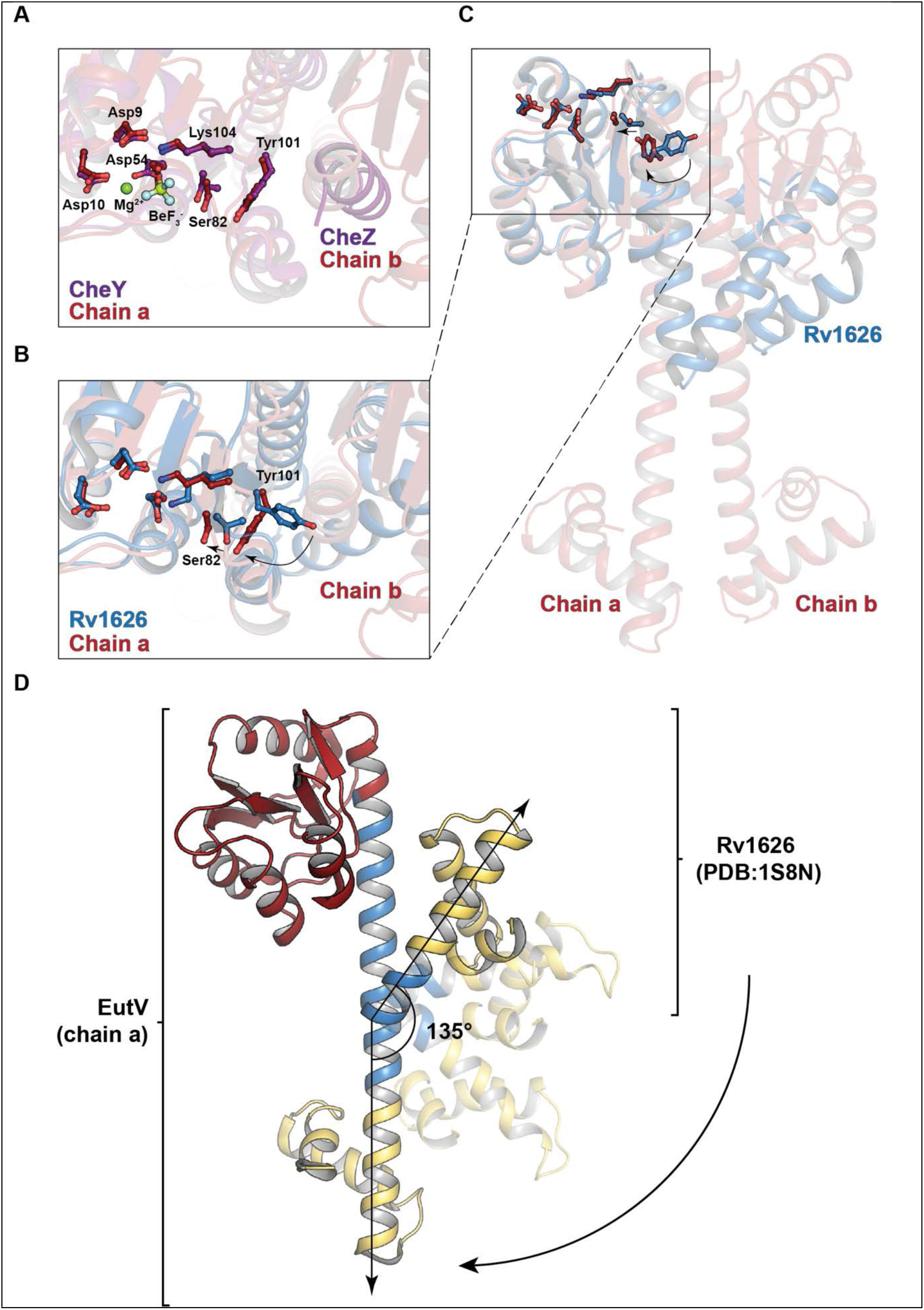
Monomer to dimer transition. (A) Chain a of the dimeric EutV structure overlaid with BeF_3_^-^ activated CheY (pdb: 2FMK) indicates the conserved residues involved in the phosphor-relay pathway are in the phosphor-activated orientation. (B-C) Overlay of the REC domain of chain a of EutV with the corresponding residues of the monomeric Rv1626 (pdb: 1s8n). Tyr101 is buried within the REC domain of EutV (relative to Rv1626) which allows chain b to associate with chain a through the α4-β5- α5 interface1. Similar dimerization is sterically blocked in Rv1626. (D) Modelled extension of the α5- α6 helix required to transition EutV from a from a monomer to a dimer. Rv1626 (pdb: 1s8n) was used to model the EutV monomer seen in solution.

A surprising revelation from the RNA bound structure of EutV was the orientation of the RNA hexaloops that contact each ANTAR domain. The RNA hexaloops face towards each other (Figure 2D, Supplementary Figure S7) and this orientation prevents the EutV dimer from simultaneously contacting both hexaloops from a single RNA molecule (Supplementary Figure S7). This positioning of the hexaloops is clearly biologically relevant given the conserved nature of the residues interacting with the RNA (Figure 4A) and the decrease in binding affinity seen when these residues are mutated to alanine (Figure 4B). Interestingly, Met172 is essential for RNA binding (Figure 4B), serving to correctly position the hexaloop on α7 by acting as a hydrophobic plug (Figure 3F). It may also act as a size determinant for the RNA loops that EutV is able to bind. Given its hydrophobic nature and proximity to Tyr164, Met172 may act to prevent smaller and more common RNA loops, such as tetraloops, from erroneously binding EutV. Methionine residues are not typically associated with RNA:protein interactions and, to our knowledge, this is the first example of this residue being obligatory for RNA binding. This finding may represent a novel mode of protein:RNA secondary structure interaction.

Position 1 and 4 in the hexaloops of the dual hairpin binding motif are conserved as adenosine and guanine bases respectively (A1 and G4) (Figure 2A-B) (25). The X-ray crystal structure has revealed the critical role G4 plays in antitermination through π-π stacking with Tyr164 (Figure 3A-C). However, A1 does not make specific contacts with EutV in the crystal structure and therefore its conserved nature is not likely due to direct interaction with EutV but rather to assist in correct RNA folding. Hexaloop structures are capable of folding into pseudotriloops through cross-loop base pairing (60, 61). Given the requirement for a G4 of the hexaloop to flip outward to interact with EutV, it is possible position 1 is conserved as an adenosine to prevent the formation of pseudotriloops that may form if other bases were present. The Met172 hydrophobic plug may also contribute to preventing pseudotriloops from folding.

The structure of the EutV bound to RNA identified two binding sites, one at each ANTAR domain of the dimer. Comparison of the REC domains and the ANTAR domains of each chain in both the RNA free and bound structures indicate no change in the secondary structure of these domains upon RNA binding (Supplementary Figure S15). Nevertheless, there is a break in symmetry of the homodimer upon RNA binding due to a larger flex in the coiled-coil of chain b, between Ile139-Glu141, than for chain a (Supplementary Figure S10). This asymmetric flex facilitates the interaction between the ANTAR domains of both chain a and chain b and the P2 hexaloop and prevents the ANTAR domain of chain A forming the same set of interactions with the P1 hexaloop. Given that the RNA bridges between ASUs and makes a crystal contact with a neighbouring ANTAR domain (Supplementary Figure S7), it is possible that this asymmetric flexing is a crystallisation requirement and does not represent a biologically relevant interaction. However, this is unlikely for two reasons: first, in the absence of any movement in chain a, the RNA-binding residues on α6-α7 of the chain b ANTAR domain would be occluded (Supplementary Figure S16). Second, the conserved nature of the residues (Figure 4A) that are positioned to bind RNA, as a consequence of the large flex in chain b, suggest a biological importance that was confirmed by alanine mutagenesis and SPR (Figure 4A-C, Supplementary Figure S12). Of particular interest was the 5-fold and 8-fold reduction in affinity of the K143A/K147A and R142A constructs respectively. Due to the asymmetric flex, these protein:RNA interactions can only occur between chain b and P2, not chain a and P1. In summary, both chains of the dimer are unable to make the same set of interactions with the RNA hairpins at the same time and furthermore, given the RNA binding orientation, there is no plausible way both hairpins of the dual hexaloop motif can contact the dimer simultaneously. This raises the possibility that the protein, in its biological role, does not contact both hairpins of the dual hairpin motif at the same time.

Within the crystal, EutV contacts two RNA hexaloops indicating the presence of two, albeit different, RNA binding sites. These hexaloops do not come from the same RNA molecule and raises the possibility that during transcription, a EutV dimer may contact two RNA hairpins from the nascent RNA of two separately transcribing RNA polymerases. This scenario is unlikely, given the rapid timescale of transcription: once the T-loop has folded, it cannot be remodelled and transcription will terminate (2). SPR studies conducted on alanine mutants confirmed that each RNA binding residue of EutV identified between the ANTAR domain of chain b and the P2 hexaloop (Figure 3A-B) play a role in binding RNA *in vitro* (Figure 4B). Upon RNA binding, the asymmetric flex of the coiled-coil domains of EutV results in a differing set of protein:RNA interactions between chains. Given it is unlikely for the EutV dimer to bind two hexaloops in *trans* and the inability of the dimer to bind both hexaloops simultaneously, it is suggested that the second RNA binding site (between the ANTAR domain of chain a and P1)(Figure 3C) provides a snapshot of a transitional binding state between a single ANTAR domain (of the dimer) and an RNA hexaloop that likely occurs prior to the full protein:RNA interactions seen between both ANTAR domains and a single hexaloop (Figure 3 A-B) (Supplementary Movie 1). This role in the initial hexaloop binding is supported by evidence that a truncated monomeric EutV (aa 132-190) retains the ability to bind a dual hexaloop motif, albeit with lower affinity than the full length (25).

If the EutV dimer only binds a single RNA hairpin, the question still remains: why are both hairpins required for antitermination? Furthermore, why is dimerization a conserved mechanism if both ANTAR domains do not make similar contact with both RNA hexaloop simultaneously (25, 35)? It is plausible that EutV dimers bind independently at each hexaloop however, as the upstream hairpin (P1) does not overlap with the T-loop, this scenario is incompatible with the obligatory requirement for both hexaloops to be present for EutV mediated antitermination *in* vivo (25). Additionally, shortening or extending the nucleotide linker between the two hexaloops, beyond the range that exists within the *eut* operon (Supplementary Figure 3B) inhibits antitermination indicating that a spatial constraint exists for the inter-loop distance (25). The possibility that binding of one EutV dimer to the upstream hairpin results in a remodelling of the P2 hairpin that facilitates a rapid binding of a second EutV dimer cannot be excluded. Similarly, the proximity of the 5’- and 3’-ends of the RNA hexaloops in neighbouring ASUs within the crystal lattice may indicate a biologically important EutV tetramer complex containing two EutV dimers and a single dual hexaloop RNA motif (Supplementary Figure S17). However, the only interface (846 Å^2^) within a tetramer complex lies between the REC domain of one EutV dimer (chain a) and the ANTAR domain of another EutV dimer (also chain a) suggesting such a binding mode would not be conserved in ANTAR proteins lacking a REC domain (Supplementary Figure S17 B). Furthermore, the residues involved in this interface have been implicated in either the correct folding of the ANTAR domain three-helical bundle or the phosphorylation of the REC domain, indicating their conserved nature may be a result of these functions, and not the formation of a larger complex (Supplementary Figure S17 C)(23, 62).

We propose a revised model for EutV mediated antitermination that most reasonably fits our observations, and those present in the literature. In the absence of ethanolamine (EA), EutV remains unphosphorylated and monomeric, leading to intrinsic termination of transcription at each T-loop (Figure 6 A-B). When present, EA stimulates EutW phosphorylation of EutV, resulting in dimerization as described in (25,34,35). Dimeric EutV binds the P1, or “recruitment”, hairpin first (Figure 6C), bringing it into proximity to the transcribing RNAP. This initial contact, as shown by the interaction between chain a and P1 in the crystal structure, is followed by a large flex in the same ANTAR domain that allows K143/K147 of the second ANTAR domain to bind to the same hexaloop, as represented by the interaction between chain b and P2 in our structure (Supplementary Movie 1). This places the second unbound ANTAR of the EutV dimer in close proximity to the RNA exit tunnel of the transcribing polymerase. As the P2, or “antitermination” hairpin, is transcribed it folds in proximity of the EutV dimer bound to the recruitment hairpin, and likely facilitates cycling of the same EutV dimer from the recruitment hairpin to the antitermination hairpin (Figure 6 C-D). Providing the EutV dimer stabilises the antitermination hairpin long enough for the polymerase to by-pass the poly-U tract, transcription will continue unabated. The recruitment hairpin may be bound by a second EutV dimer, after the first dimer has transitioned to the antiterminator hairpin, which would likely prevent a backwards transition of a EutV dimer. This model provides the most rational explanation for both the inability of the EutV dimer to contact both hexaloops from a single RNA molecule, as seen in our crystal structure and the requirement for both hairpins to exist in order to achieve *in vivo* antitermination (25) . Furthermore, both ANTAR domains of the EutV dimer are utilised, although not simultaneously, providing the rationale for the dependence on EutW, and thereby EA-induced EutV phosphorylation/dimerization, for antitermination *in vivo* (25). Finally, this model explains the spatial constraint that is applied to the linker between hairpins of the dual hexaloop motifs of the *eut* operon (25).

**Figure 6.**
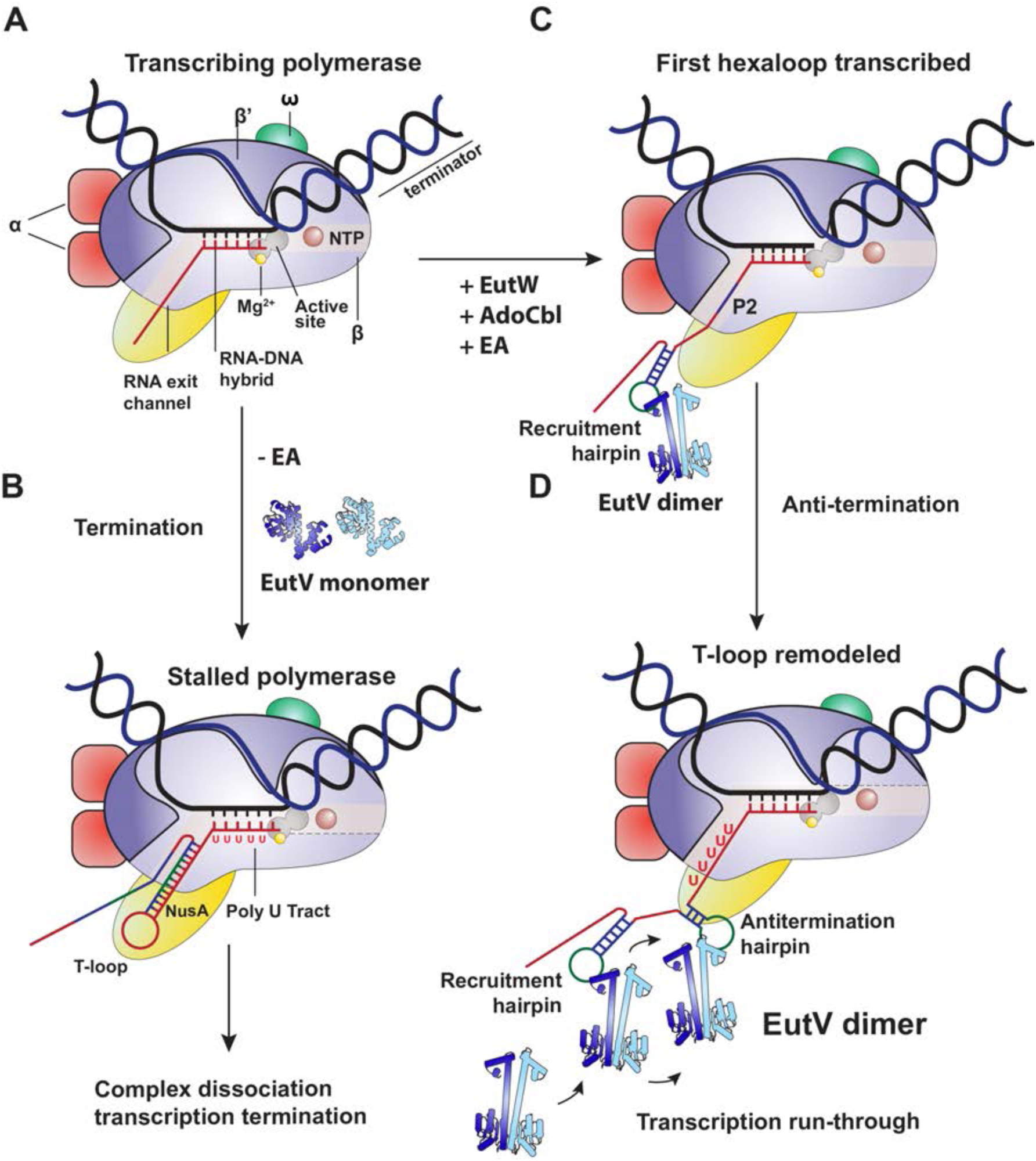
Proposed model for EutV antitermination. (A) Core RNA polymerase (RNAP) (in bacteria composed of an α-dimer, a β subunit, a β′ subunit and an ω subunit) is bound to the DNA duplex composed of the template strand (black) and the non-template strand (blue), and the nascent RNA (red). (B) Stalled RNAP on poly-U tract. Intrinsic T-loop formation within the RNA exit tunnel is stabilised by NusA (yellow). (C) In the presence of EutW, AdoCbl and ethanolamine (D) dimeric EutV binds the first hexaloop (P1) of the dual hairpin motif before cycling to the (D) second hexaloop (P2) as transcription continues.

As dimeric EutV is unable to bind both hexaloop simultaneously we have redefined the minimal ANTAR domain target motif to a single hexaloop motif. This allows for potential single hexaloop ANTAR binding sites to exist within the bacterial genome when the time constraint of a transcribing polymerase is absent. Indeed in *E. faecalis* this may be true for the sequestration of EutV by the small non-coding *eutX* RNA (*rli55* in *L. monocytogenes* (63)) in the absence of essential cofactors required for EA metabolism (Supplementary Figure S18). Inspection of the *eutX* sequence identified an additional single hexaloop, with a G4 nucleotide, in addition to the previously described dual hexaloop motif (64). Furthermore, recent work describing the presence of RNA stemloops that overlap with ribosome binding sites and 5’-UTR of transcripts in *Mycobacterium tuberculosis* do not always obey the classical dual hexaloop motif (37). Further bioinformatic analysis and experimental work is required to determine to what degree single hexaloop motifs play in ANTAR domain function and what novel processes they may regulate.

## MATERIAL AND METHODS

### Protein expression and purification

The gene sequence for full length EutV from *E. faecalis* was purchased as a synthetic gene block and cloned into an isopropyl-β-D-thiogalactopyranosid (IPTG)-inducible expression vector containing a TEV protease cleavable his-tag at the N-terminus. *E. coli* Rosetta™ 2(DE3)pLysS cells were transformed with plasmid DNA and were grown overnight on LB agar plates containing 25 μg mL^-1^ Kan and 17 μg mL^-1^ Cam. Large scale cultures, inoculated with resuspended colonies from the transformation plate, were grown at 37°C until an OD_600_ of between 0.3-0.4 was reached. The growth temperature was lowered to 18°C and cultures were grown to an OD_600_ of 0.6 when cells were induced with 0.5 mM IPTG and subsequently grown for a further 16-20 hours before harvested via by centrifugation.

EutV was purified via a multi-step chromatography protocol consisting of immobilised nickel ion affinity chromatography, proteolytic cleavage of the his-tag, anion exchange chromatography and finally size exclusion chromatography. Briefly, bacterial cells were lysed by sonication in 25 mM sodium phosphate pH 7, 1 M NaCl, 10% w/v glycerol, 1 mM TCEP, 5 mM imidazole, 1× cOmplete EDTA-free protease inhibitor, 10 μg mL^-1^ DNase I and 100 μg mL^-1^ lysozyme. The lysate was clarified via centrifugation and the supernatant was applied to a 5 mL HisTrap FF column (GE) to immobilise his-tagged proteins. Bound proteins were washed with buffers containing increasing concentrations of imidazole before final elution with a buffer containing 500 mM imidazole. Proteolytic cleavage of the his-tag using TEV protease was performed in conjunction with overnight dialysis into 25 mM sodium phosphate pH 7, 150 mM NaCl, 10% w/v glycerol and 1 mM TCEP in preparation for a second round of immobilised nickel chromatography to remove the cleaved his-tag. To remove contaminating nucleic acids, the sample was then subjected to anion exchange chromatography using a 1 mL ResourceQ column (GE) equilibrated with dialysis buffer, where EutV remained unbound to the column whilst contaminating nucleic acids and proteins were retained. EutV was subsequently concentrated and gel filtration was performed using a Superdex75 column (GE) into a final size exclusion buffer containing 50 mM HEPES pH 7, 300 mM NaCl, 10% w/v glycerol and 1 mM TCEP.

### *In vitro* transcription

For crystallographic studies, the 51-nt *eutP* RNA was produced by *in vitro* transcription. A linearized DNA template containing the *eutP* RNA sequence, a 5’ T7 RNA polymerase protomer site and a 3’ HDV ribozyme was incubated with T7 RNA polymerase (100 μg mL^-1^, produced in house), pyrophosphatase (10 μg mL^-1^) and RiboSafe RNAse inhibitor (20 U mL^-1^) in transcription buffer (50 mM HEPES pH 7.5, 40 mM MgCl_2_, 100 μg mL^-1^ BSA, 2 mM spermidine, 40 mM DTT and 7.5 mM of each NTP) for 2 hours at 37°C before the reaction was finalised at 42°C for 2 hours. To ensure maximum folding and subsequent cleavage of the 3’ HDV ribozyme, 50 μg mL^-1^ of an 18-bp oligo (complementary to the 3’ end of the RNA) was added post-transcription to prevent secondary RNA structures from inhibiting cleavage. The RNA mixture was then subjected to 2 min heating at 95°C, cooling to 53°C for a further 2 min, followed by rapid cooling on ice for 5 minutes. This process was repeated 2-4 times. The *eutP* RNA was separated from the template plasmid via denaturing polyacrylamide gel electrophoresis (38) and excised from the gel before extraction by dilution in MQW.

### Electromobility shift assay (EMSA)

#### Preparation of ^32^P labelled nucleic acid probes

All ^32^P RNA used for EMSA experiments were generated by 5’ end labelling. Purified RNA constructs (200 pmol) were 5’ labelled in 20 μL reactions by incubation (37°C, 1 hour) with γ-^32^P ATP (10 mCi mL ^-1^) and 5 U T4 polynucleotide kinase (T4 PNK) as per the manufacturer’s instructions. Reactions were ended by the addition of EDTA (50 μM) and unincorporated γ-^32^P ATP removed by centrifugation using a Mini Quick Spin RNA column (1,000 x *g*, 4 minutes). RNA was diluted to 200 μL with SDS extraction buffer (1% SDS, 50 mM EDTA pH 8, 100 mM tris-HCl pH 7.5), and subsequently precipitated by the addition of ammonium acetate (0.3 M, pH 5.2) and 3 volumes of cold ethanol (−20°C) and incubated on dry ice (1 hour). The precipitated RNA was pelleted by centrifugation (15,000 x *g,* 10 minutes) and washed with 70% (v/v) cold ethanol before being resuspended in sterile MQW and stored at -20°C.

#### Native PAGE

6% (29:1 acrylamide/bis) gels were cast in 1x TBE buffer supplemented with 2.5 mM MgCl_2_ and were pre-run at 4°C for 2-3 hours (200 V, 50 mA, amperage limiting). Varying concentrations of protein (0-100 μM) were incubated with a constant trace quantity of ^32^P-labelled RNA probe in gel shift buffer (50 mM HEPES pH 7, 150 mM NaCl, 10% glycerol, 0.016% [w/v] bromophenol blue and 0.04% [w/v] xylene cyanol) for 1 hour on ice prior to running on the gel. Gels were electrophoresed for 4 hours (200 V, 50 mA, amperage limiting) at room temperature using an ASG-250 adjustable slab gel kit (C.B.S Scientific, Thermo Fisher Scientific) before being transferred to Whatman 3 MM paper and dried (50°C, 2 hours) using a gel dryer (Model 583, BioRad) attached to a vacuum pump. Dried gels were exposed to a phosphoscreen for 16-24 hours and imaged directly using a Typhoon FLA900 (GE Healthcare).

### Purifying RNA bound EutV

EutV was incubated with fresh BeF_3_ buffer (30 mM NaF, 5 mM BeSO_4_ and 2.5 mM MgCl_2_) for 1 hour. Precipitation was removed by centrifugation (5 min, 13500 x *g* and 4°C) before the sample was added to *eutP* RNA at a three molar excess and incubated for 1 hour. Assembled complexes were purified from their individual components by gel filtration in complex-forming buffer (25 mM HEPES pH 7, 150 mM NaCl, 1 mM TCEP, 2.5 mM MgCl_2_ and 2.5% w/v glycerol).

### Size exclusion chromatography (SEC) multi-angle laser light scattering (MALS)

Experiments were performed on an ÄKTA FPLC system (GE Healthcare) equipped with miniDAWN TREOS MALLS and Optilab T-rEX detectors (Wyatt Technology). Purified EutV was separated using a Superdex200 10/300 Increase (GE Healthcare) in EutV size exclusion buffer with UV absorbance monitored at 215, 260 and 280 nm. Astra software (Wyatt Technology) was used for data analysis, including baseline and peak broadening corrections using a dn/dc value of 0.1852 mL g^-1^.

### Nuclear magnetic resonance (NMR) spectroscopy

Protein samples were prepared in size exclusion buffer supplemented with D_2_O (10% (v/v)) and 2,2- dimethyl-2-silapentane-5sulfonic acid (DSS, 150-300 μM). Samples were placed in 3 mm NMR tubes (Shigemi) and spectra were acquired using Bruker Avance III 600 or 800 MHz NMR spectrometers, each fitted with a cryogenic TCI probe-head, at 4°C. Spectra were processed using TOPSPIN (Bruker, Karlsruhe, Germany) and ^1^H chemical shifts were directly referenced to DSS at 0 ppm.

### Far-UV circular dichroism (CD) spectroscopy

Purified protein was dialysed (overnight at 4°C) into 5 mM HEPES pH 7, 300 mM NaF and 1 mM TCEP using 10 kDa molecular weight cut-off dialysis tubing (Progen, Darra, QLD) and diluted to 5-15 μM. CD spectra were recorded using a Jasco J-185 spectrometer (ATA Scientific) at 4°C using a 1 mm quartz cuvette (Sigma-Aldrich). Data were recorded using a speed of 20 nm min^-1^, a response time of 1 s, and a sensitivity of 20 mdeg over the wavelength range 250-195 nm. Data are the average of three scans and were buffer baseline-corrected.

### Surface plasmon resonance

SPR measurements were made using a Biacore T200 instrument (GE Healthcare). Experiments were performed at 4 °C using a multicycle kinetic titration method. 3’ biotinylated RNA constructs (linked via an extended TEG spacer arm) were purchased from Integrated DNA technologies (Baulkham Hills, NSW) and immobilised on a Biotin CAPture chip (GE Healthcare) in 10 mM sodium acetate pH 4.8, 150 mM NaCl, 2.5 mM MgCl_2_ and 0.05% Tween, with a target density of ∼200-250 RU, as per manufacturer’s protocol. Different concentrations of proteins were flowed over the reference and RNA-immobilised cells at a flow rate of 50 µL min^-1^ using 10 mM HEPES, pH 7, 150 mM NaCl, 0.05% Tween-20 and 2.5 mM MgCl_2_ as the running buffer. Stripping and regeneration of the chip surface was performed as per the manufacturer’s instructions using the supplied reagents. All data were analysed and fit to a 1:1 Langmuir binding isotherm using the Biacore T200 Evaluation Software.

### Statistical Analysis

The affinity of EutV to each of the three RNAs was compared using an analysis of variance (ANOVA), with the individual means being compared using a Tukey’s HSD test to maintain an overall 5% error rate. The overall p value was 0.06 and none of the means were different at the 5% level.

### X-ray crystallography

EutV crystals used for diffraction studies were crystallised in 0.075 M tris pH 8.5, 18.75% v/v tert-Butanol and 25% v/v glycerol using a sitting-drop vapour-diffusion method at 18 °C, with crystals taking between 4-10 days to form. The EutV:RNA complex was crystallised in 0.1 M NaCl, 0.1 M HEPES pH 7.5 and 1.4-1.8 M ammonium sulphate with crystals taking 14 days to reach full growth. Crystals were cryoprotected using 25% glycerol and frozen by plunge-freezing in liquid nitrogen. X-ray diffraction data were collected from frozen crystals at the Australian Synchrotron using the Macromolecular Crystallography MX2 beamline (microfocus) at 100 K and a wavelength of 0.9537 Å (39). XDS was used to integrate data and the data were processed further using the CCP4i suite (40, 41). Indexing, scaling, and merging of the data was performed using AIMLESS (42, 43). Initial phases were calculated by molecular replacement using PHASER (44). The REC domain of Rv1626 (30) was modelled as a dimer using AmiR (19) (PDB: 1S8N and 1QO0 respectively) and used as an initial search model. The models were visualised in COOT (45) and were built manually by iterative rounds of refinement using phenix.REFINE (44) and ISOLDE (46) until convergence. MOLPROBITY (47) was used for structure validation and identification of steric clashes and geometric problems in the final model. Surfaces were evaluated using the web based PISA software (48). The quality of the final models were validated using the wwPDB server and submitted to the PDB (6WSH and 6WW6 for EutV alone and RNA bound respectively). Structure diagrams were generated using PyMOL. The data collection and refinement statistics for these structures are outlined in Table S1.

## ACCESSION NUMBERS

Atomic coordinates and structure factors for the EutV crystal structures have been deposited with the Protein Data bank under accession numbers 6WSH (EutV alone) and 6WW6 (EutV:RNA bound) and will be released upon publication.

## ACKNOWLEDGEMENTS

This research was undertaken using MX1 and MX2 beamlines at the Australian Synchrotron, part of ANSTO, and made use of the Australian Cancer Research Foundation (ACRF) detector. We also thank Dr Santosh Panjikar and Prof. Mitchell Guss for the positive discussion and advice during structure determination, Dr Alastair Stewart and Yi Zeng for the critical reading of the manuscript, Adrienne Kirby from the University of Sydney NHMRC Clinical Trials Centre for assistance with statistical analysis and The Bosch Institute for access to SPR instrumentation.

---

**Supplementary Figure S1.**
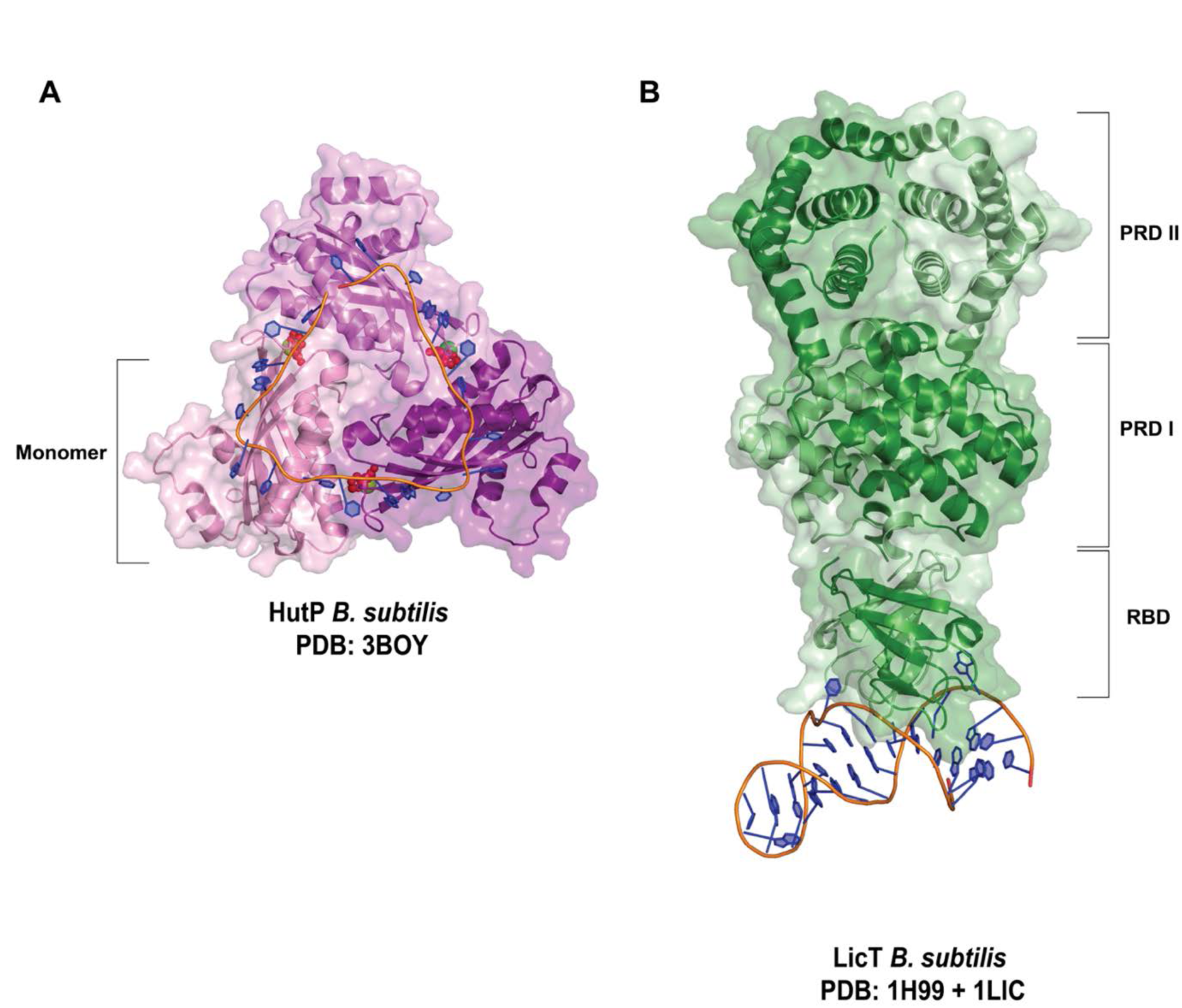
(A) HutP (*B. subtilis*) hexamer bound to its cognate RNA sequence (PDB: 3BOY). Histidine ligands, shown as small red spheres, and Mg^2+^ ions, as large green spheres, bind at the trimer interfaces, supporting biochemical evidence that proper HutP function requires these ligands. NAG triplets bind to each monomer. (B) Modelled complex of N-terminal LicT (*B. subtilis*) bound to RNA (PDB:1LIC) and C-terminal LicT PRD domains (*B. subtilis*) (PDB 1H99). Phosphorylation events in the PTS regulation domain (PRD) II and PRD I dictate dimerization of LicT, which facilitates binding a structured RNA element through the RNA binding domains (RBD). PyMOL cartoon and surface representation of three characterized RNA binding antiterminators. Ligands are presented as small red spheres, RNA bases as blue rings and the phosphate backbone as an orange tube.

**Supplementary Figure S2.**
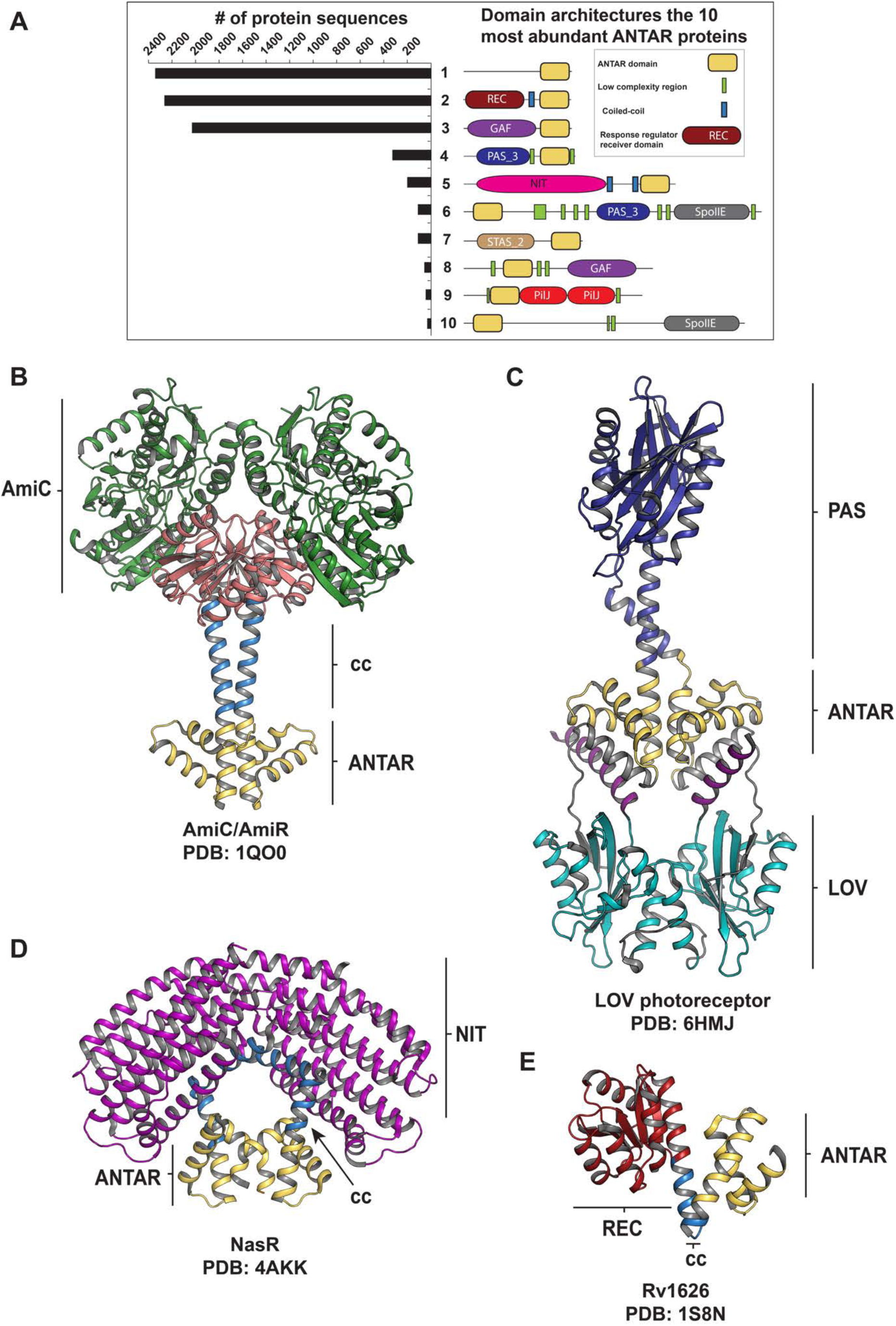
Structures of known ANTAR domain proteins and distribution of ANTAR domain proteins according to their domain organization (A) Bar graph showing the number of ANTAR proteins that contain the domain organisation shown schematically. Figure adapted and updated from (1). (B-E) ANTAR domains colored yellow and predicted coiled-coil domains in sky-blue. All structures are in an inactive state. (B) The *P. aeruginosa* amidase operon anti-terminator AmiR in complex with the negative regulator AmiC (green) (PDB 1QO0). AmiR forms an intimate dimer through a coiled-coil domain and pseudo-receiver domain (salmon). (C) Light-oxygen-voltage (LOV) photoreceptor PAL from *N. multipartite.* PAS and LOV domains colored in blue and cyan respectively (D) nasREDCBA operon anti-terminator protein NasR from *K. oxytoca* (PDB 4AKK). Nitrate/nitrite sensing (NIT) domain colored in purple. (E) Putative anti-terminator protein Rv1626 from *M. tuberculosis* (PDB 1S8N). Two component response regulator (RR) domain colored in red.

**Supplementary Figure S3.**
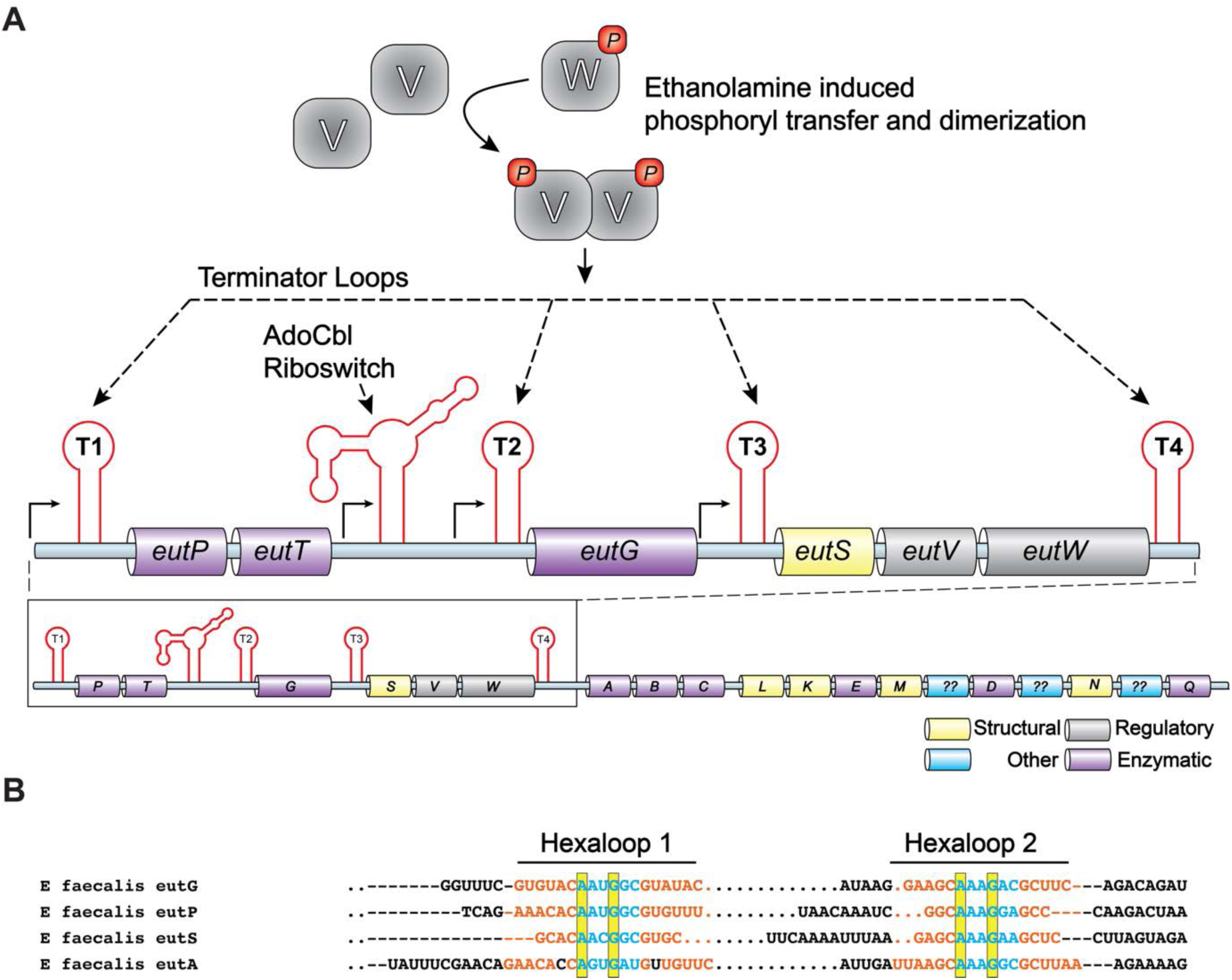
Schematic representation of genes and regulatory elements within the *eut* operon from *E. faecalis*. Genes within the *eut* operon allow for the efficient catabolism of ethanolamine and provide host bacteria with a source of carbon and/or nitrogen. Histidine kinase EutW phosphorylates EutV in the presence of ethanolamine inducing dimerization (A) Four intrinsic terminator loops and a riboswitch (shown in red) facilitate the regulation of expression of genes within the *eut* operon. Dimeric EutV disrupts the formation of the intrinsic terminator loops. (B) Primary sequence of the four dual hexaloop anti-terminator elements from the *eut* operon. Bases involved in stem formation are shown in orange, hexaloop bases in blue and conserved bases at position 1 and 4 of the hexaloops are highlighted in the yellow box.

**Supplementary Figure S4.**
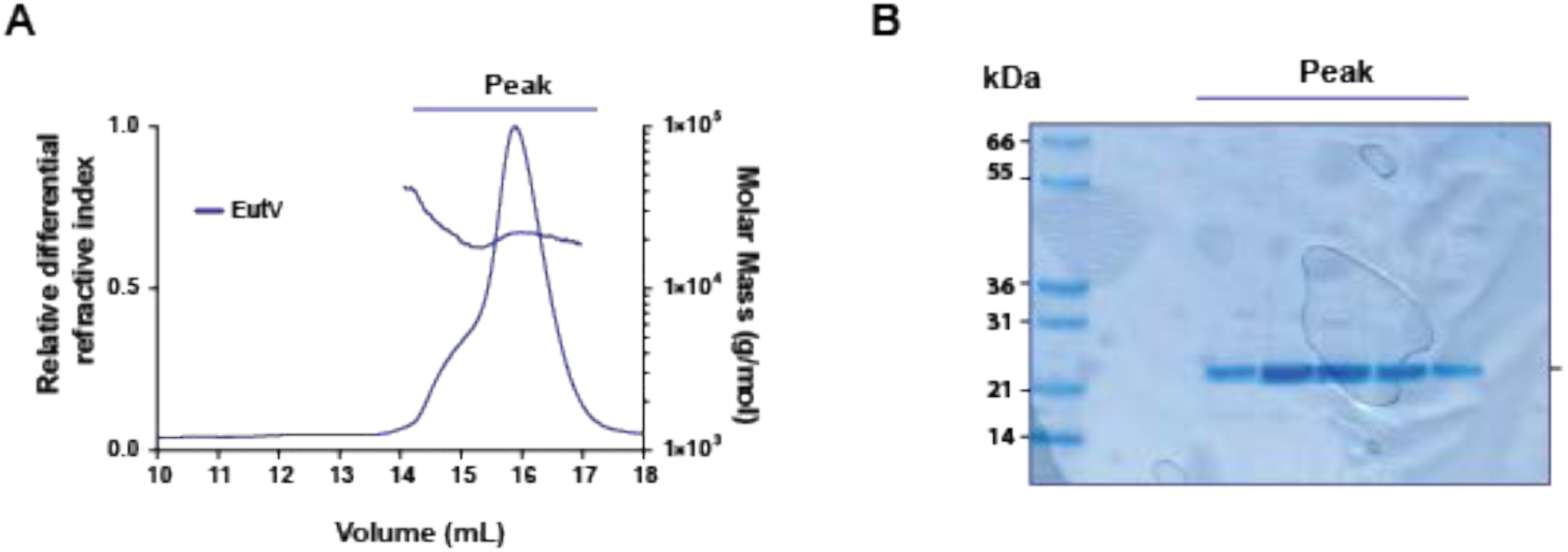
SEC-MALS light scattering analysis of EutV. (A) Size exclusion chromatogram showing differential refractive index on the left y-axis as a relative measure of concentration for EutV. Molecular mass estimates are displayed as a scatter plot over each peak (right-hand y-axis). EutV was separated on a Superdex200 10/300 Increase column and MALS analysis of the peak reveals the protein is largely monomeric (21.9 ± 0.6 kDa) in solution, although a larger molecular weight species is also present. (B) SDS-PAGE showing the higher molecular weight species is not caused by a contaminating protein, suggesting that a concentration dependant dimer is present.

**Supplementary Figure S5.**
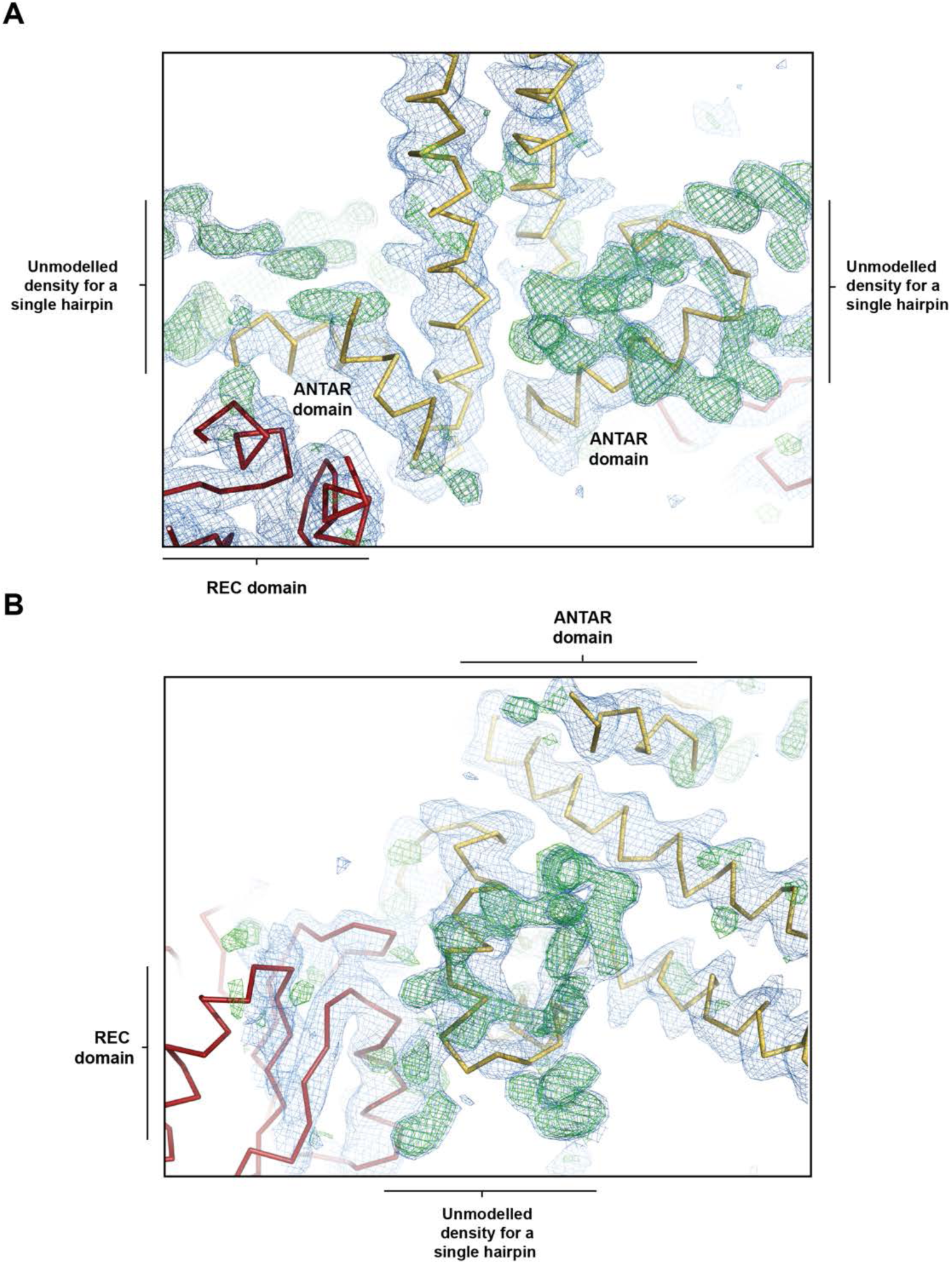
Unmodeled electron density for RNA hairpins. Electron density map of EutV:EutP dataset after molecular replacement and rigid body refinement using the CheY-like and ANTAR domains of EuV. 2F_o_-F_c_ and F_o_-F_c_ maps are contoured to at 1.5 σ and 3 σ respectively. The N-terminal CheY-like domain and C-terminal coiled coil/ANTAR domains are shown in red and yellow respectively.

**Supplementary Figure S6.**
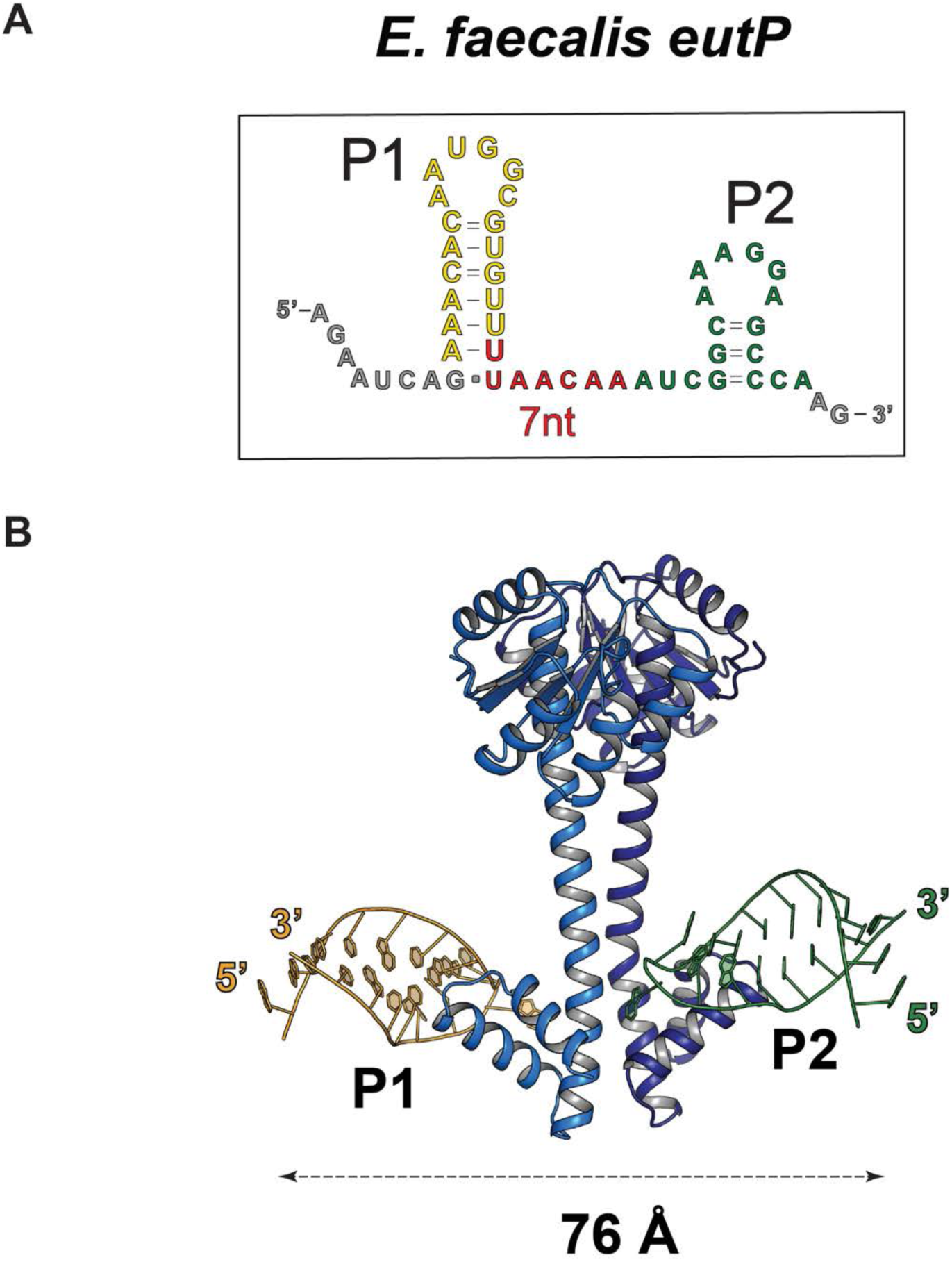
Distance between model P1 and P2 hexaloop hairpins. (A) Primary sequence of the 51-nt RNA used for co-crystallisation with EutV. Yellow and green bases represent the successfully modelled P1 and P2 loops respectively. Bases in grey represent the unmodeled 5’ and 3’ ends and in red for the unmodeled linker. (B) Cartoon representation showing the modelled P1 and P2 loops with the distance between the 5’ end of each indicated.

**Supplementary Figure S7.**
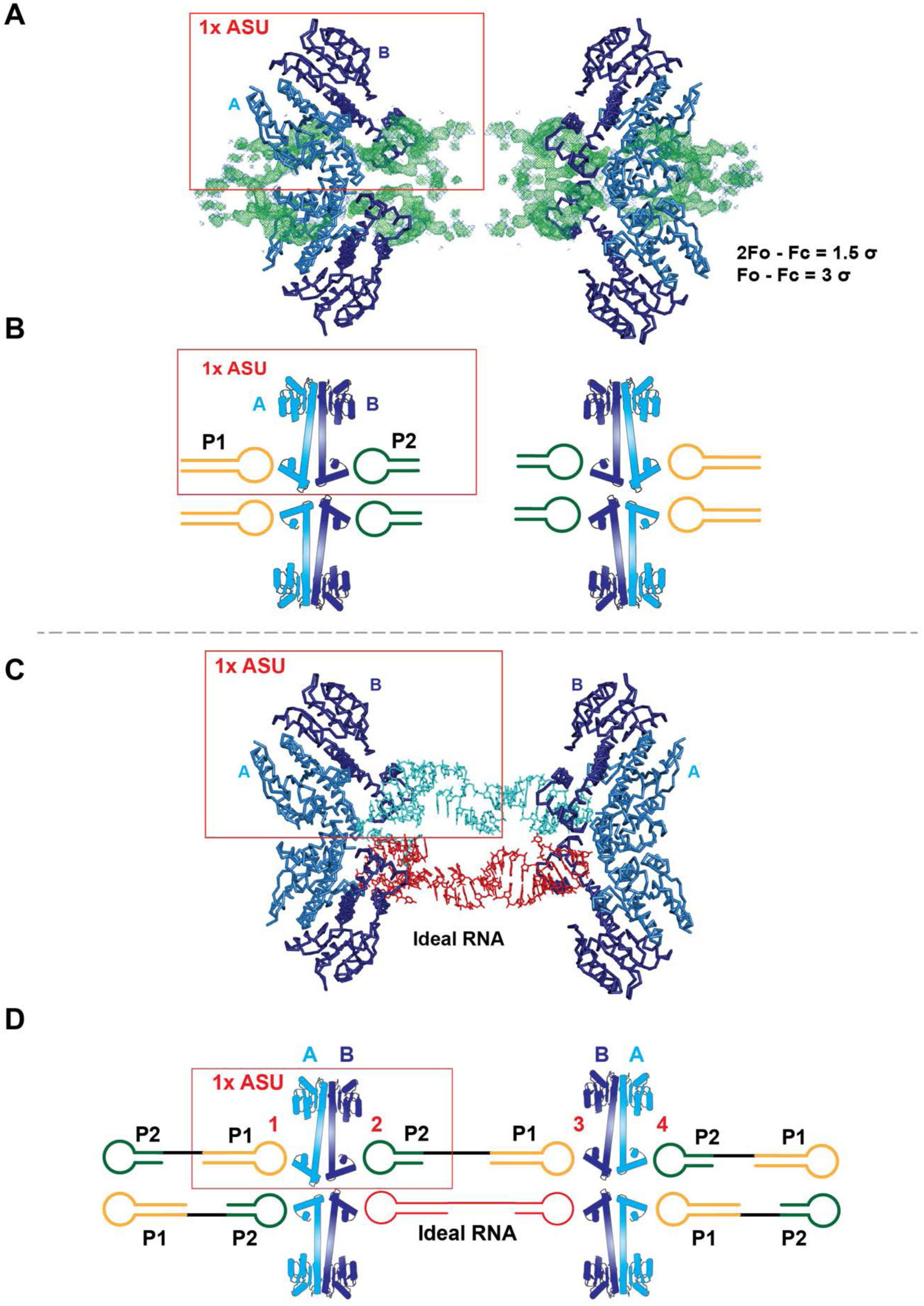
Crystal packing of the RNA bound structure. Each asymmetric unit (red box) is composed of one dimer of EutV and electron density for two RNA hairpins. Four asymmetric units (ASU) are shown in (A), the orientation of the hairpins relative to the EutV chains is shown schematically in (B). (C) Idealised *euP* RNA model (cyan and red) manually fitted between two ASUs. RNA generated with *RNAComposer* (2). (D) Schematic diagram of the EutV:*eutP* RNA crystal lattice. The four possible orientations of EutP RNA contacting the EutV dimer are numbered in red. Chain A and B of the protein is coloured in cyan and dark blue respectively. P1 and P2 hexaloops are coloured in green and cyan respectively. 2F_o_-F_c_ and F_o_-F_c_ maps contoured to 1.5 σ and 3 σ respectively.

**Supplementary Figure S8.**
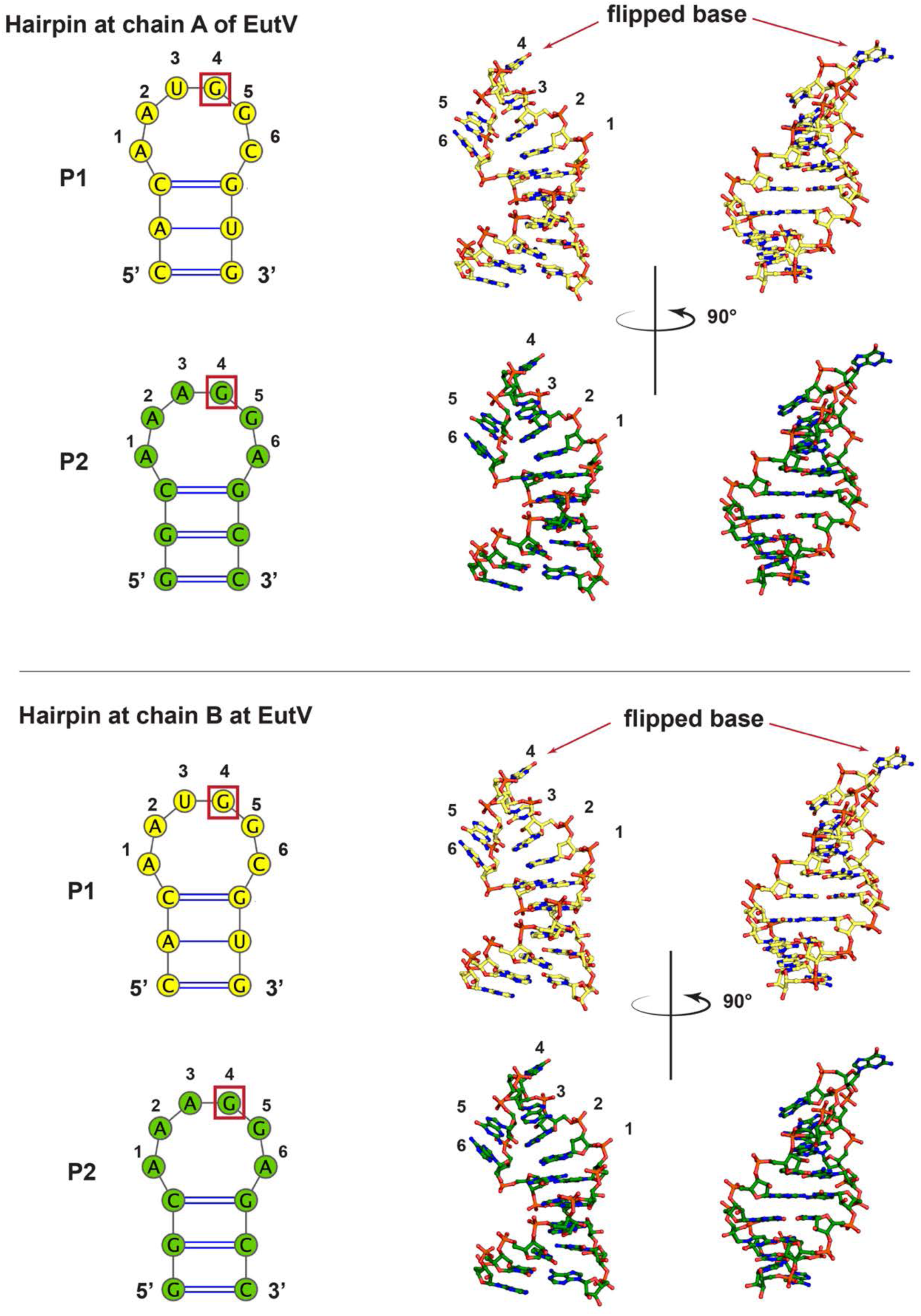
Cartoon representation of the two modelled hairpins (P1 and P2) at either ANTAR domain of the EutV dimer. After dual-occupancy refinement in *Phenix.refine* (3), both P1 and P2 hairpins and the ANTAR domain of each chain of EutV refined to near identical positions.

**Supplementary Figure S9.**
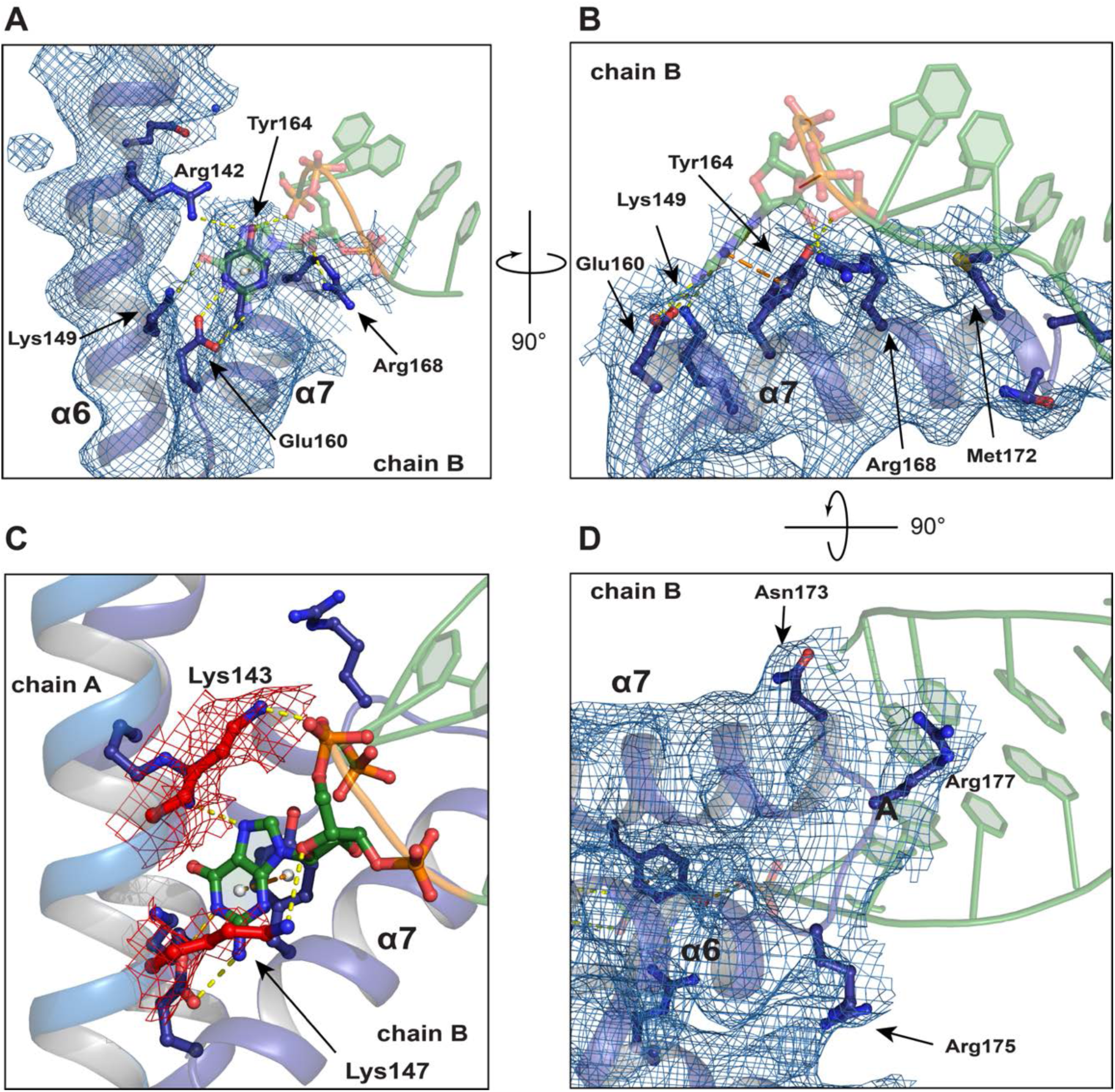
Electron density of RNA binding residues from EutV. 2F_o_-F_c_ map contoured to 1 σ and colored blue in all panels except (C) where colored red. Hydrogen bonds shown as yellow dashes. π- π stacking represented as orange dashes.

**Supplementary Figure S10.**
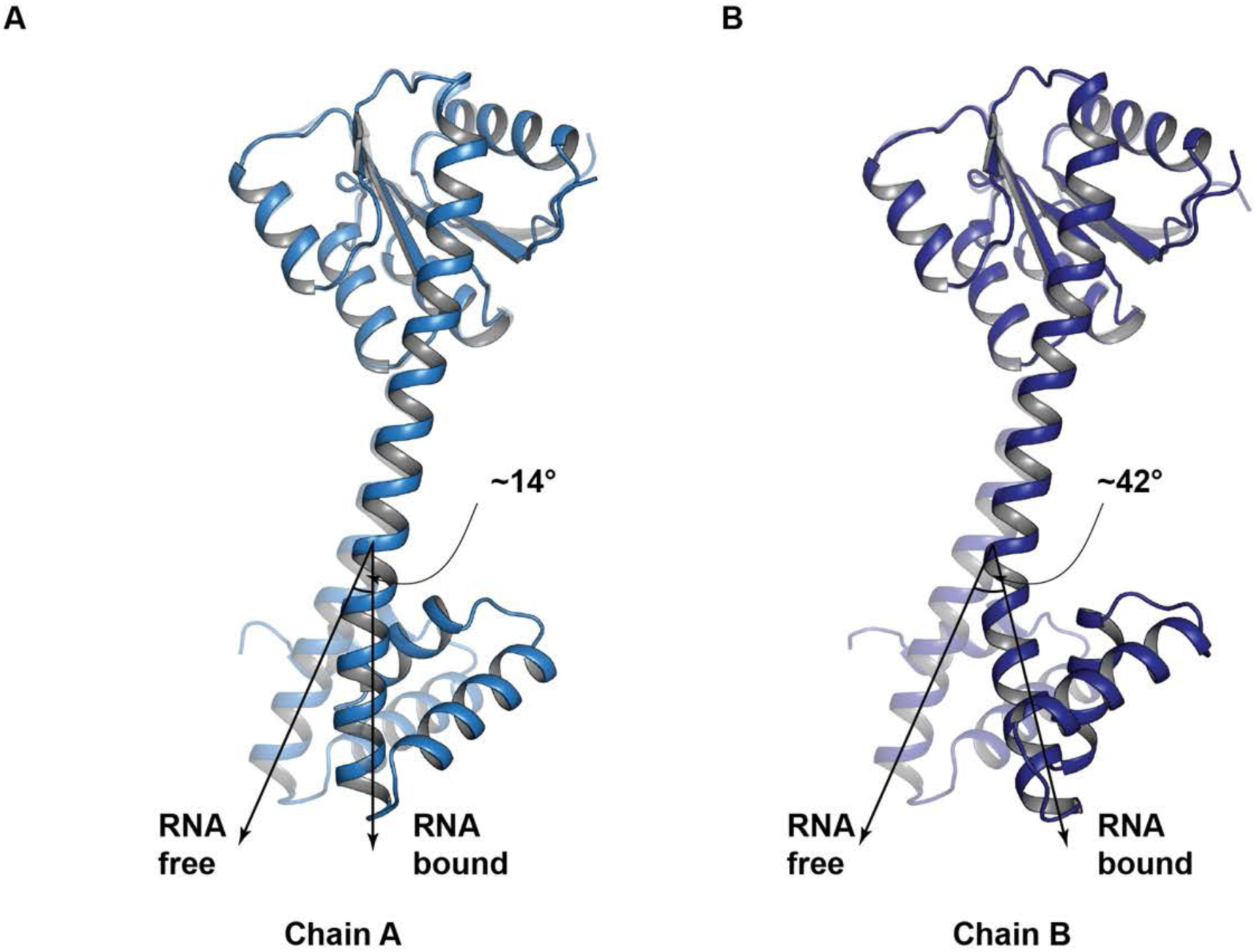
Comparison of RNA free and bound EutV dimers. Comparison between each chain of the RNA free (transparent cartoon) and RNA bound (non-transparent cartoon) EutV dimers. (A) Chain A (sky-blue cartoon) moves approximately 14° upon RNA binding while (B) chain B (dark blue cartoon) moves over 40°.

**Supplementary Figure S11.**
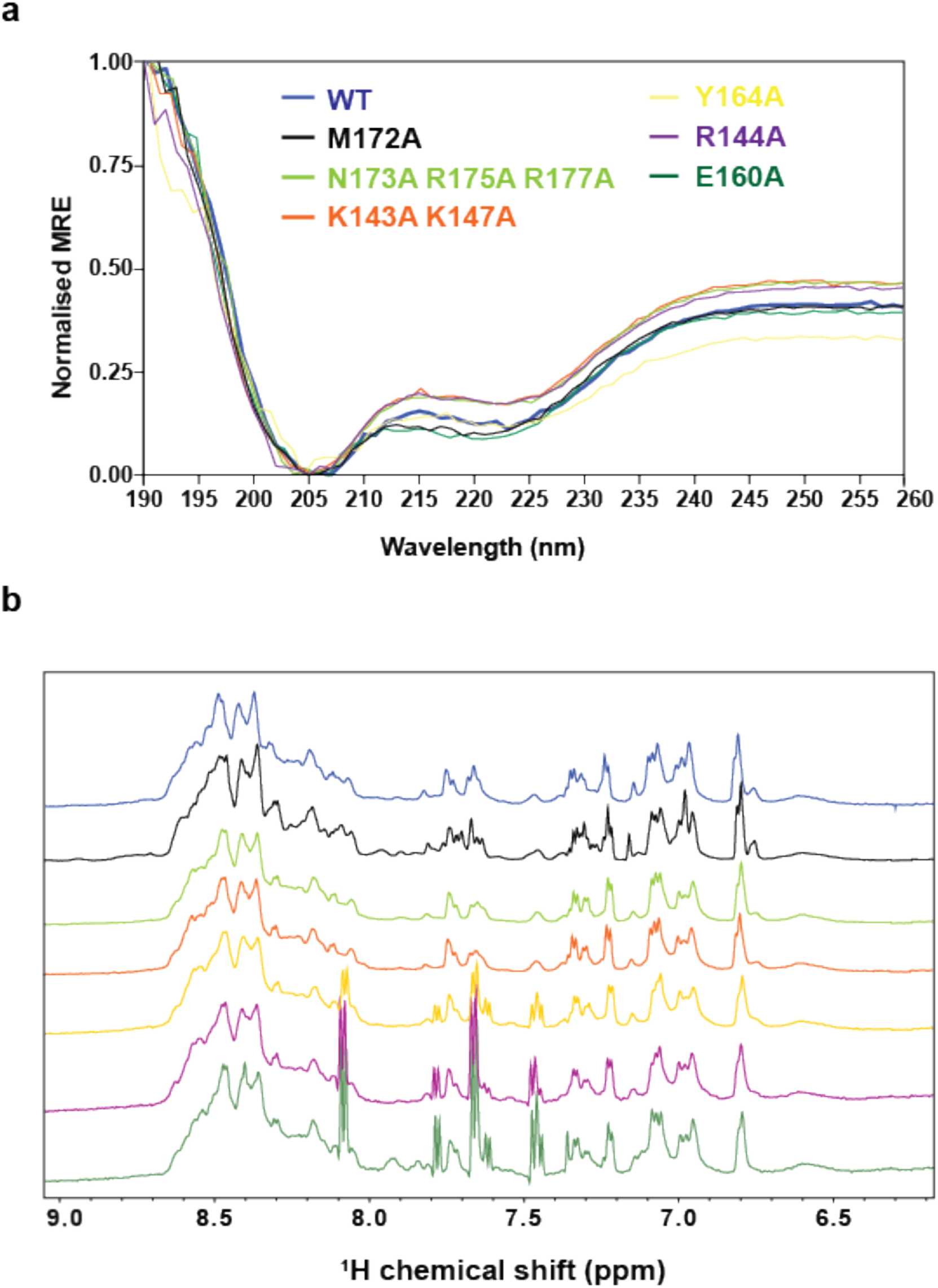
UV CD and 1D 1H NMR spectrum of wild-type and mutant EutV constructs. Coloring is consistent across all panels. (A) Far-UV CD spectra normalized by setting the mean residue ellipticity (MRE) at 205 to 0. Spectra were recorded at 4°C with ∼10 μM protein samples prepared in 10 mM HEPES pH 7, 300 mM NaF and 1 mM TCEP. Spectra are an average of three consecutive measurements. (B) 1D ^1^H NMR spectra showing peaks in the amide (>6 ppm) region, recorded at 4° with 100 μM protein samples prepared in 50 mM HEPE3S pH 7, 300 mM NaCl, 1 mM TCEP.

**Supplementary Figure S12.**
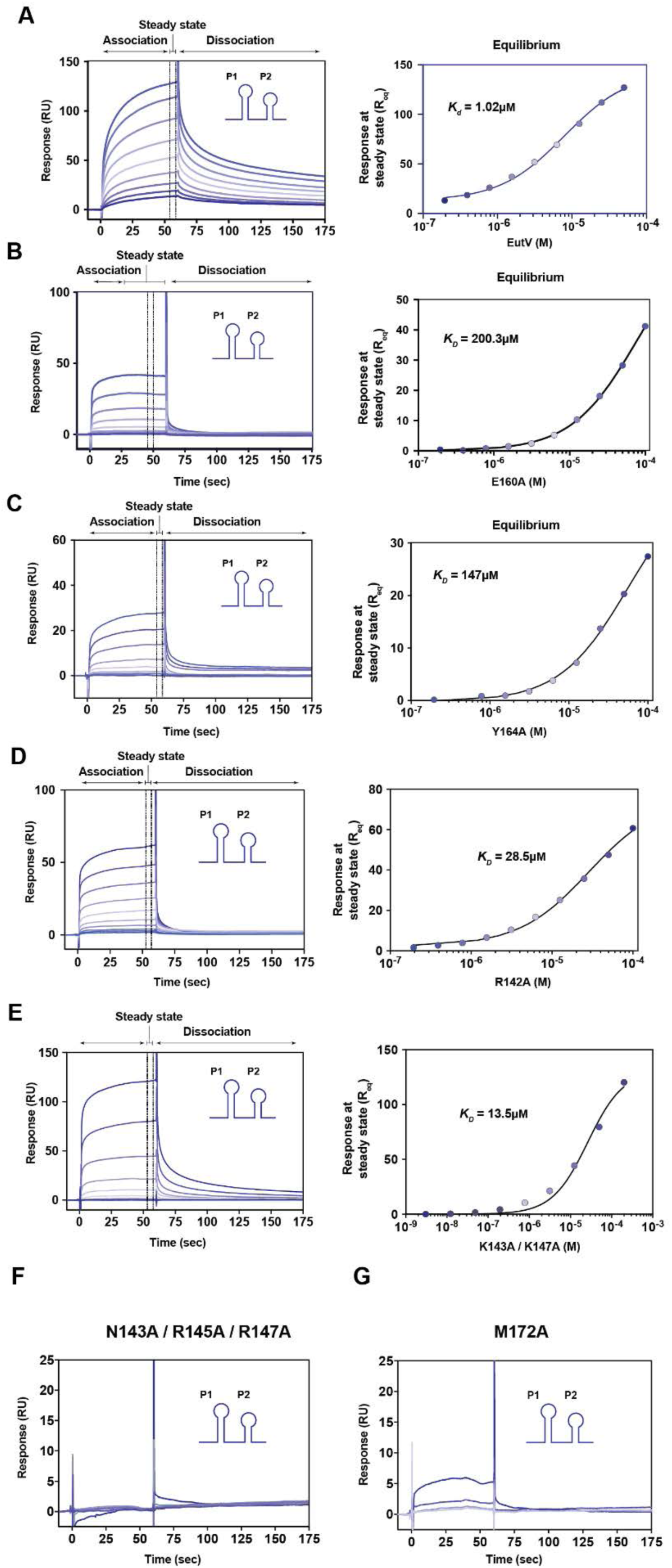
Representative SPR experiments of EutV WT and mutant constructs binding EutP RNA. ((A-E) Left panels) Representative normalised SPR sensorgrams of EutV WT and mutant proteins binding to EutP RNA. Association and dissociation regions are shown in the top most panel and apply for all sensorgrams. Sensorgrams show ten increasing EutV concentrations (0, 0.4, 0.8, 1.5, 3.125, 6.25, 12.5, 25, 50, 100 μM). (A-E, Right panels) Representative dose response plot of the interaction of EutV with immobilised RNA (as shown in corresponding left panel) at equilibrium fitted to a one-site Langmuir isotherm. The R_max_ parameter for the fragment binding was fixed during the curve fitting process and as estimated using the fitted maximum response for the positive control EutV WT. (F-G) Representative normalised SPR sensorgram of EutV N143A / R145A / R147 and EutV M172A respectively showing no binding.

**Supplementary Figure S13.**
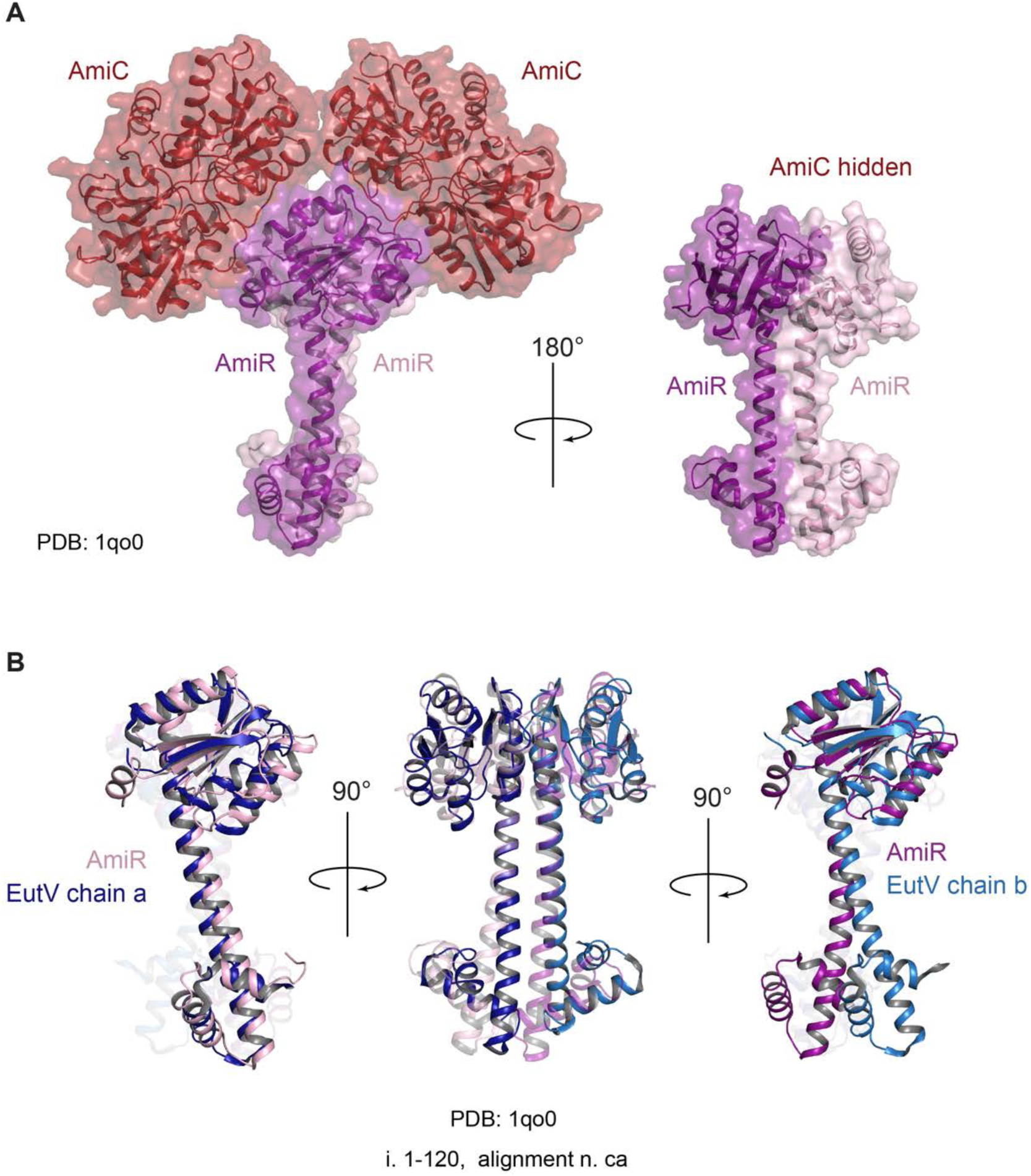
Structural alignment of EutV and AmiR. (A) Crystal structure of AmiC/AmiR complex (pdb: 1qo0) with AmiC coloured in red and AmiR in a dark and light shades of pink. (B) Structural alignment of EutV and AmiR showing the asymmetric nature of the AmiR (attributed to crystal packing limitations (4)) dimer compared to symmetric EutV dimer. EutV is coloured in a light and dark shades of blue.

**Supplementary Figure S14.**
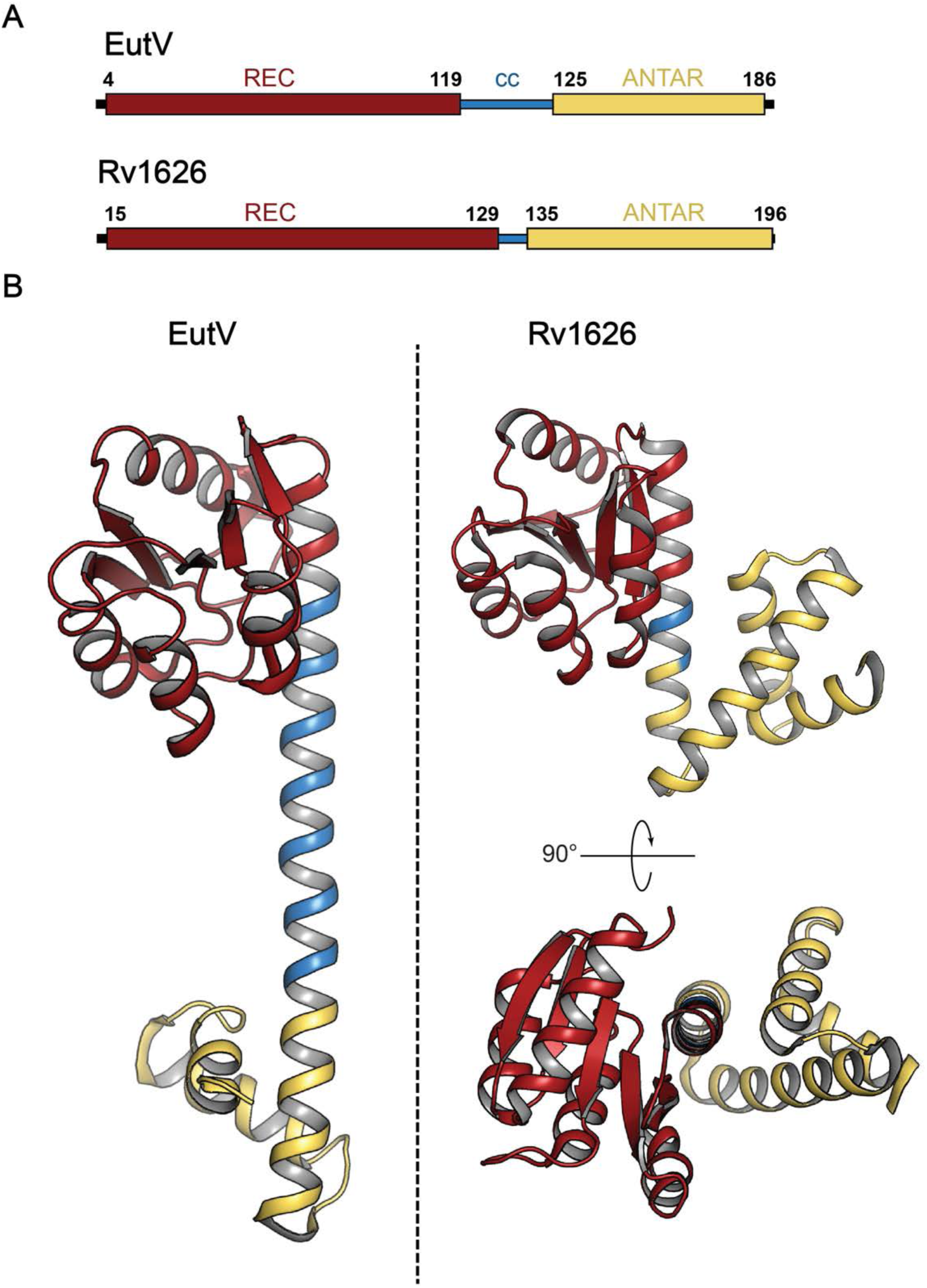
Structural comparison of EutV and Rv1626. (A) Domain organisation of EutV and the putative transcription anti-terminator Rv1626 from *M. tuberculosis* (5) (B) Comparison of a single chain of the RNA free EutV dimer and Rv1626. Both proteins are coloured the same way with the receiver (REC) domain is shown in red, the coiled-coil domain in blue and the ANTAR domain in yellow.

**Supplementary Figure S15.**
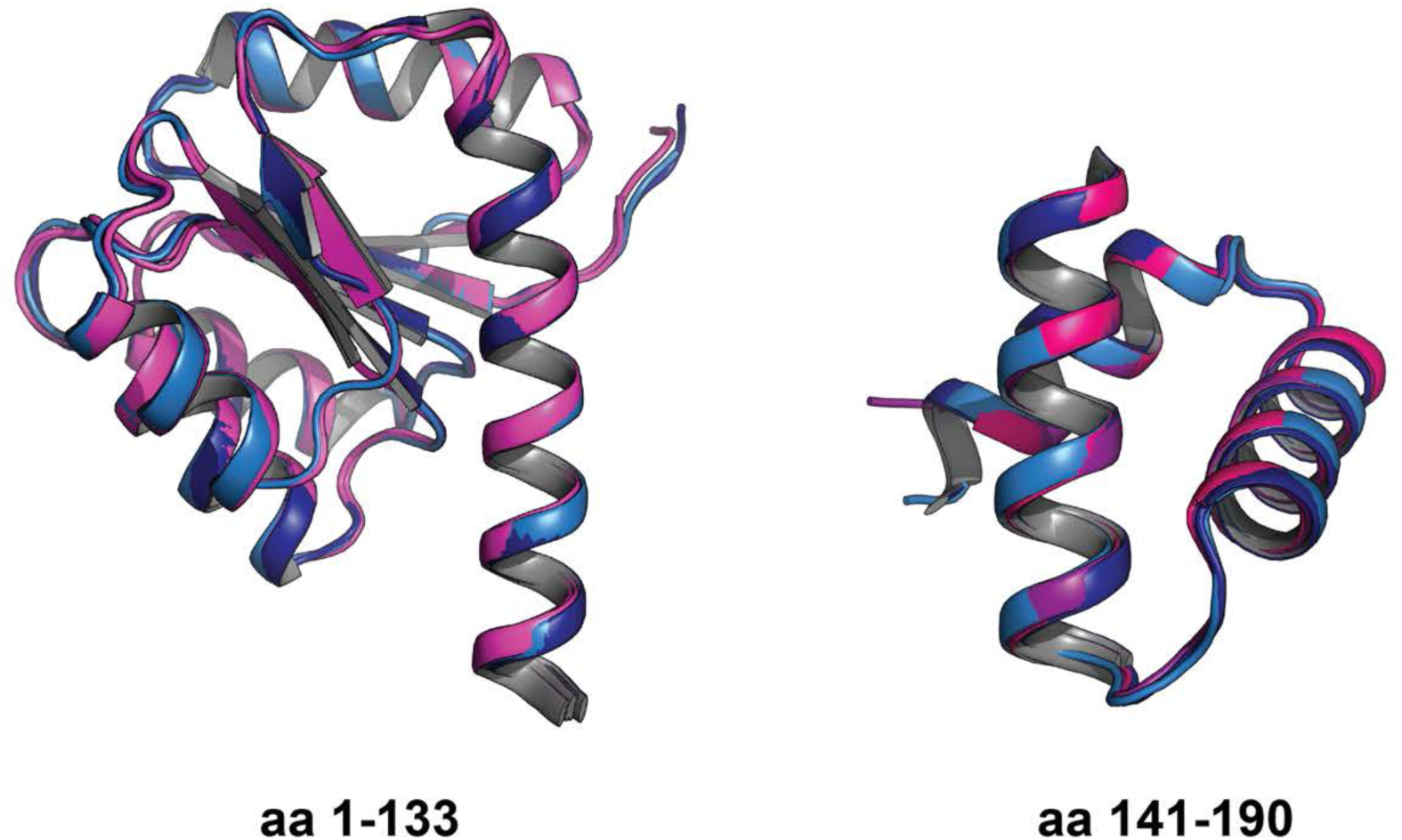
Comparison of REC domains and ANTAR domains of EutV free and the EutV:EutP RNA structure. Chain a and b of EutV:EutP RNA complex show no significant difference when compared to both chains of the RNA-free structure (RMSD Cα atoms 0.36 Å and 0.30 Å) for REC and ANTAR domains respectively. EutV RNA free is coloured in a light and dark shades of pink for chain a and b respectively while EutV:EutP RNA is coloured in a light and dark shades of blue for chain a and b respectively.

**Supplementary Figure S16.**
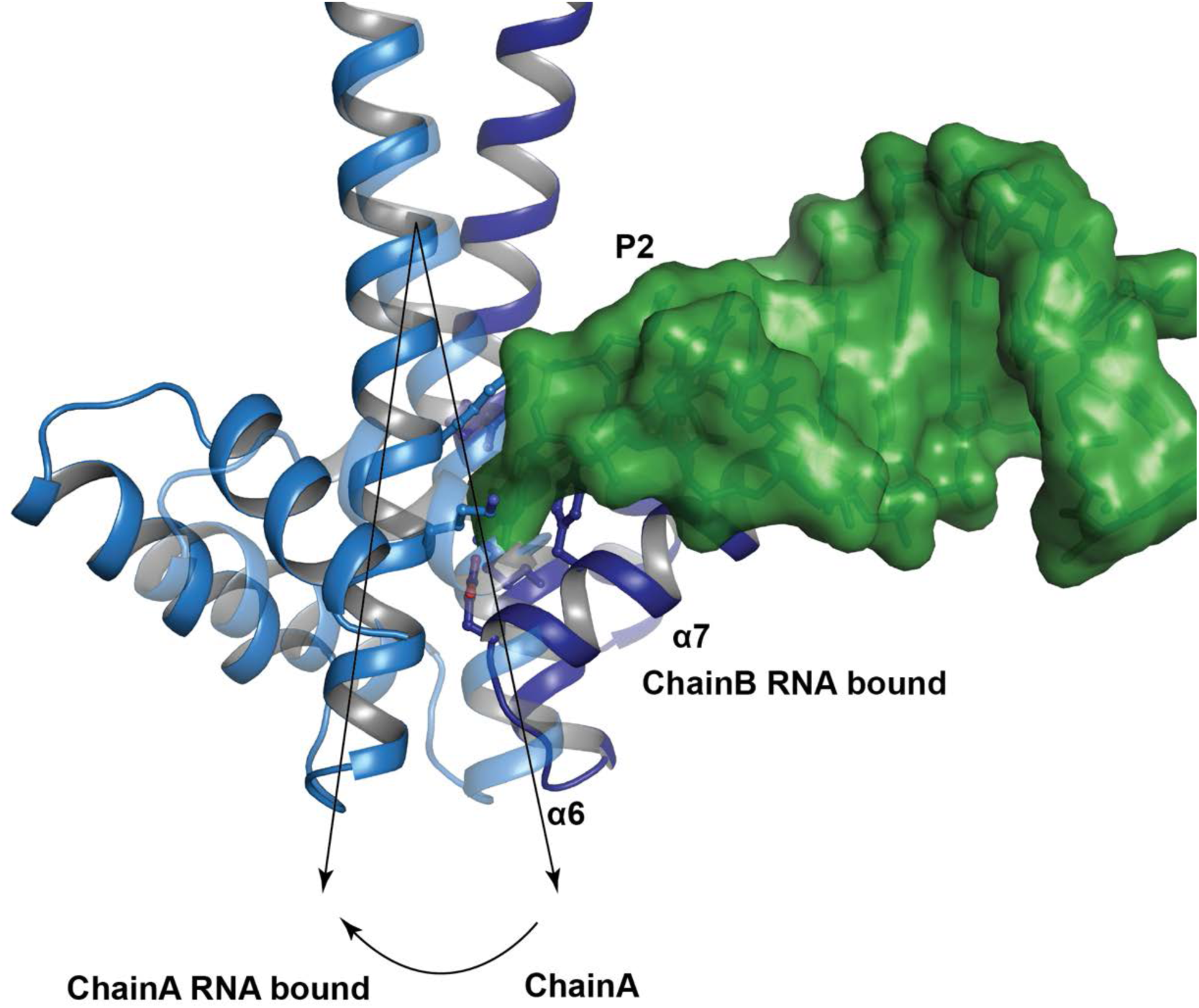
Helical flexing upon RNA binding. Cartoon showing flexing of α6 between RNA free (transparent) and RNA bound EutV structures is required to expose the RNA binding surface of α7.

**Supplementary Figure S17.**
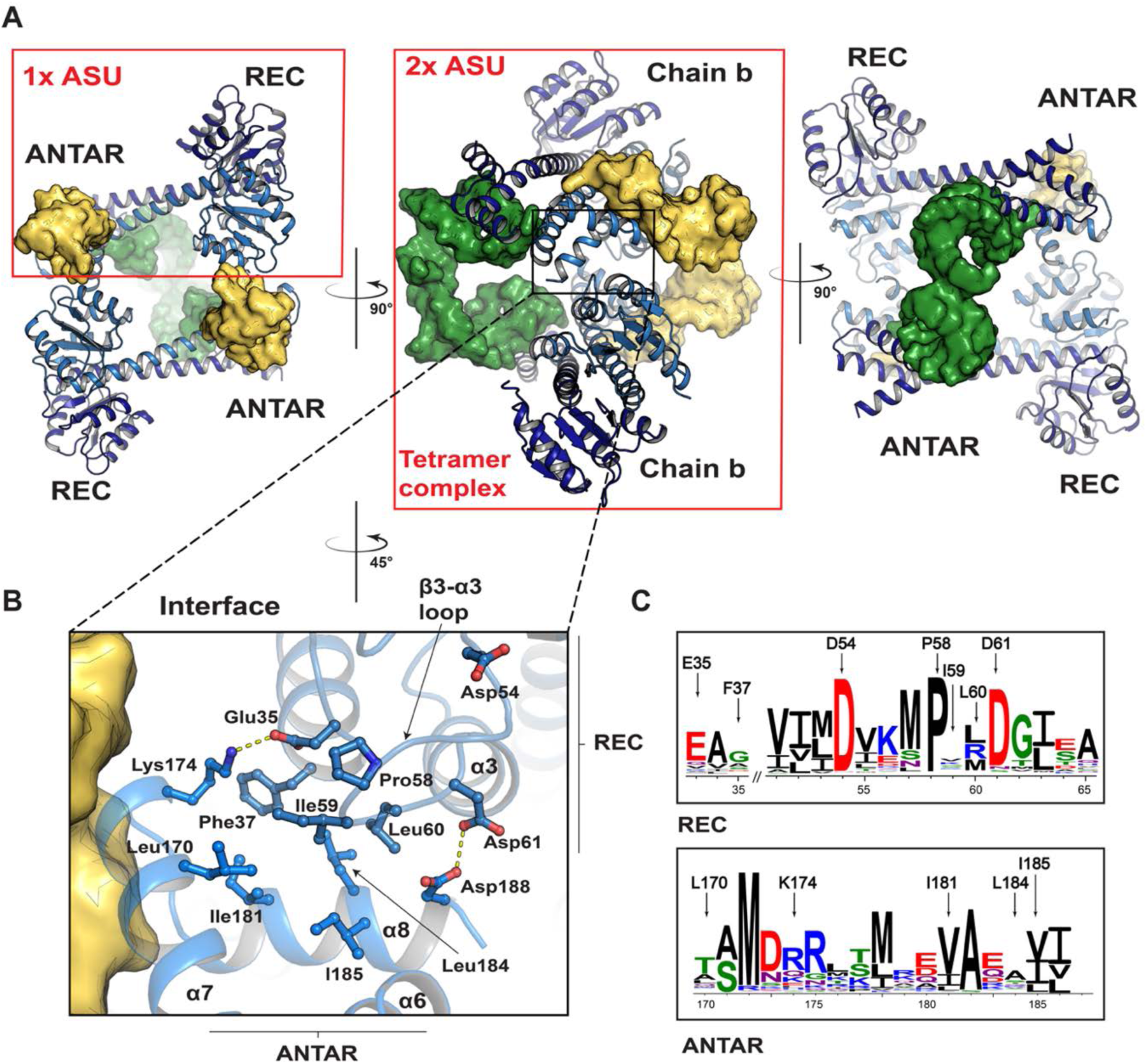
Interaction of potential EutV tetramer within crystal lattice. (A) Lattice packing arrangement of EutV tetramer complex. Two EutV dimers pack head to tail within the lattice. Symmetry related RNA shown as yellow and green surface representation. EutV chain a and b shown as sky-blue and dark blue pymol cartoon respectively. (B) Crystal contact between chain a of two symmetry related dimers (within possible tetramer complex). Hydrogen bonds represented as yellow dashes. (C)) Multiple sequence alignment 2332 ANTAR domain proteins containing a N-terminal REC domain and C-terminal ANTAR domain shown as a WebLogo (6).

**Supplementary Figure S18.**
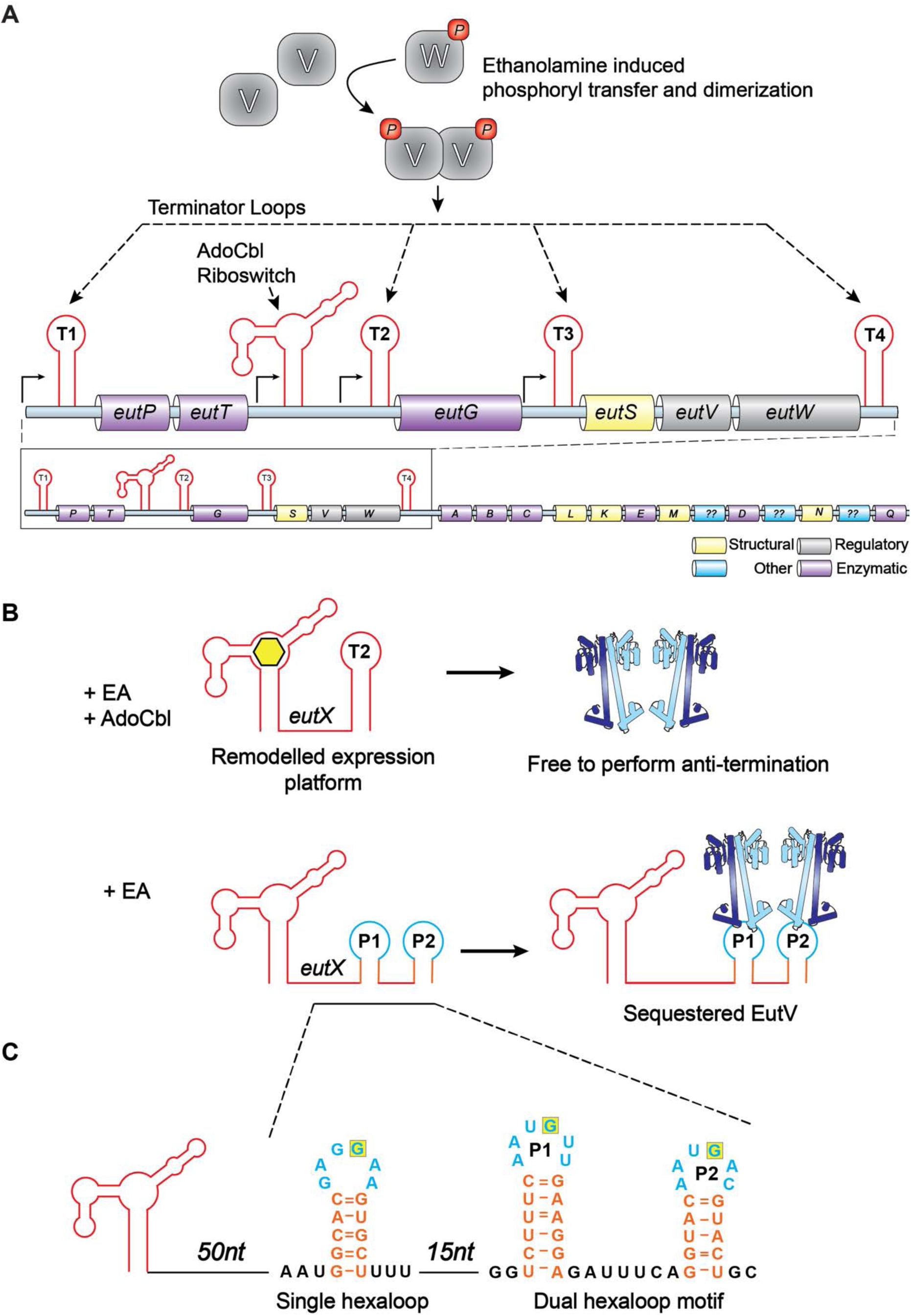
EutX sRNA regulation. (A) Gene organisation of the eut operon highlighting the eutT-eutG intragenic region (B) Model for eutX transcript regulation of the eut operon by sequestration of dimeric EutV as proposed by Deboy et al. (7) Adenosylcobalamin (AdoCbl)(yellow hexagon), and essential cofactor for ethanolamine catabolism, binds the AdoCbl riboswitch (red cartoon) remodelling the expression platform to promote T-loop formation (single red loop). When ethanolamine is present, but AdoCbl absent, dimeric EutV is sequested by a eutX transcript containing hexaloops. (C) Cartoon showing additional single hexaloop structure identified in the 131nt region between AdoCbl riboswitch and dual hairpin motif. Hairpins and hexaloops shown in orange and blue respectively. Position 4 of the hexaloop coloured in yellow. Sequence analysed using MFOLD (8).

Supplementary Movie 1: Shows the RNA binding residues, flexing of helices and the interactions with RNA between the EutV dimer crystal structure, the transition state model and RNA bound crystal structure. Morphs between states were generated in PyMOL Transitions state model is composed of chain a of the RNA free structure with chain a of the RNA bound structure.

**Supplementary Table 1:**
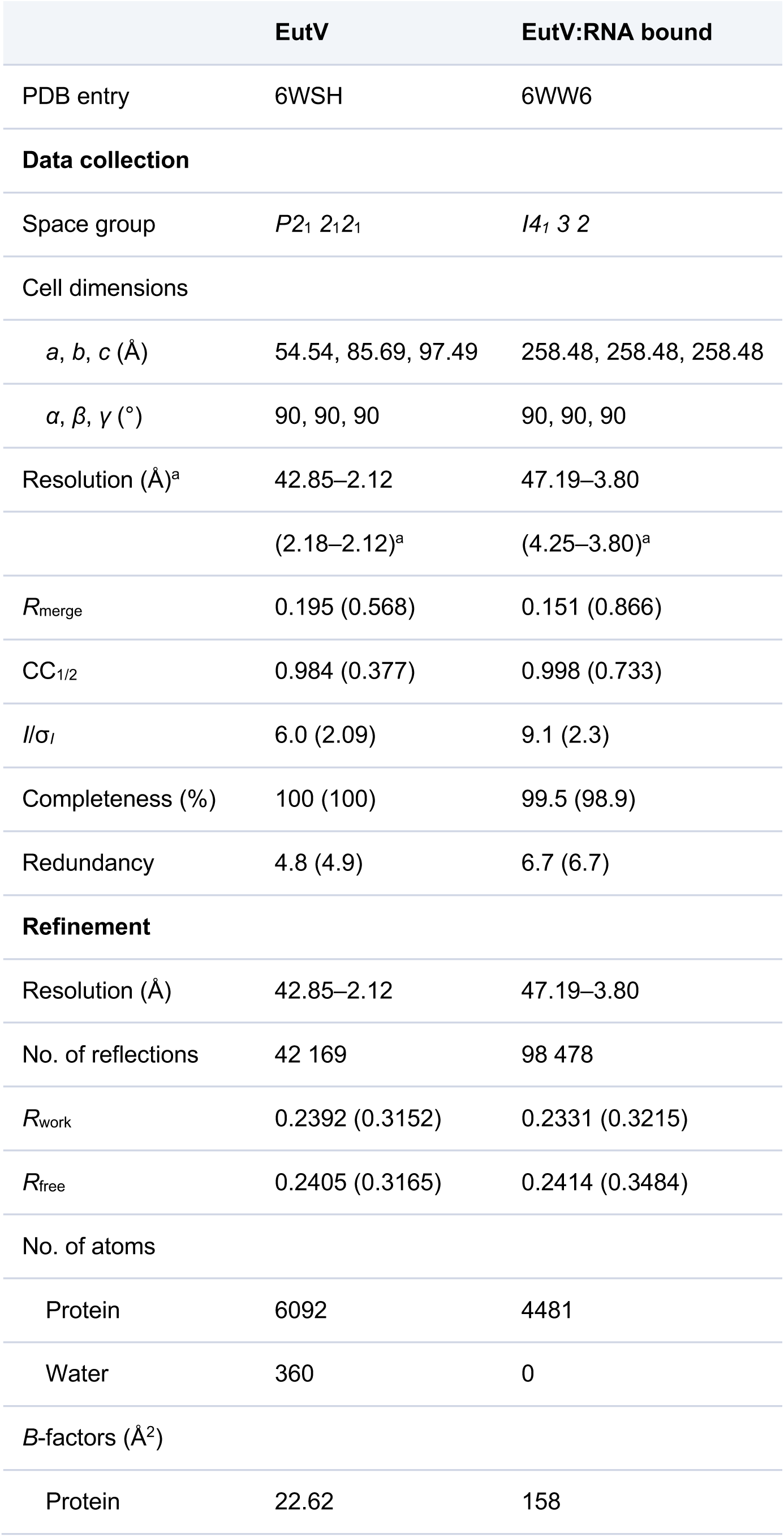
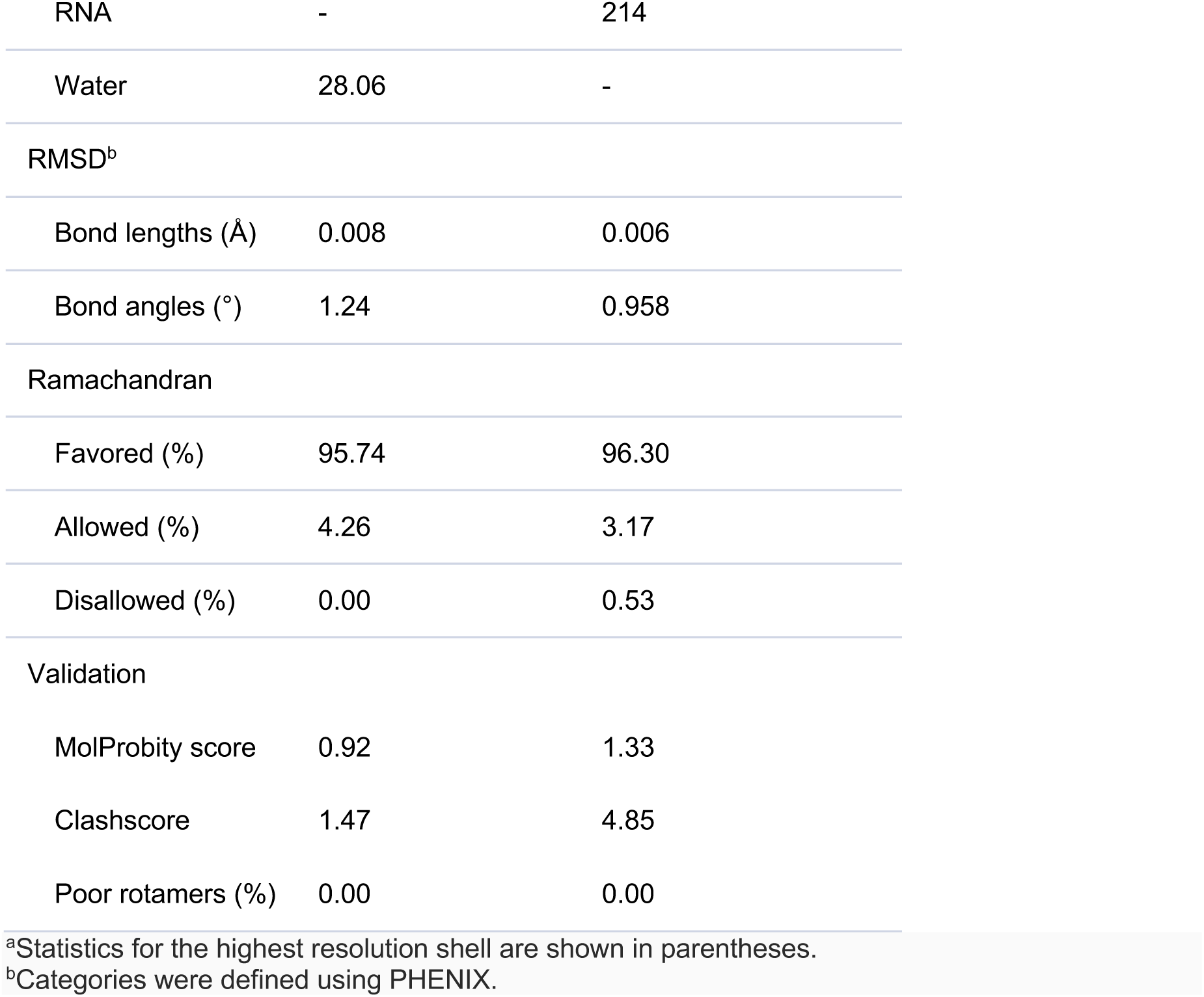
X-ray diffraction and crystal structure refinement statistics for EutV alone and RNA bound complex

